# Making sense of the linear genome, gene function and TADs

**DOI:** 10.1101/2020.09.28.316786

**Authors:** Helen S Long, Simon Greenaway, George Powell, Ann-Marie Mallon, Cecilia M Lindgren, Michelle M Simon

## Abstract

**Background:** Topologically associating domains (TADs) are thought to act as functional units in the genome. TADs co-localise genes and their regulatory elements as well as forming the unit of genome switching between active and inactive compartments. This has led to the speculation that genes which are required for similar processes may fall within the same TADs, allowing them to share regulatory programs and efficiently switch between chromatin compartments. However, evidence to link genes within TADs to the same regulatory program is limited.

**Results:** We investigated the functional similarity of genes which fall within the same TAD. To do this we developed a TAD randomisation algorithm to generate sets of “random TADs” to act as null distributions. We found that while pairs of paralogous genes are enriched in TADs overall, they are depleted in TADs with CCCTC-binding factor (CTCF) ChIP-seq peaks at both boundaries. By assessing gene constraint as a proxy for functional importance we found that genes which singly occupy a TAD have greater functional importance than genes which share a TAD, and these genes are enriched for developmental processes. We found little evidence that pairs of genes in CTCF bound TADs are more likely to be co-expressed or share functional annotations than can be explained by their linear proximity alone.

**Conclusions:** These results suggest that algorithmically defined TADs consist of two functionally different groups, those which are bound by CTCF and those which are not. We detected no association between genes sharing the same CTCF TADs and increased co-expression or functional similarity, other than that explained by linear genome proximity. We do however find that functionally important genes are more likely to fall within a TAD on their own suggesting that TADs play an important role in the insulation of these genes.

## BACKGROUND

The organisation of the mammalian genome in three dimensional space is non-random and hierarchically organised (1). Using Hi-C (2) it was shown that chromosomal loci are clustered into two, mega base scale structures known as the A and B compartments (3). The A compartment is enriched for active, euchromatin whereas the B compartment is enriched for inactive, heterochromatin (3, 4). The formation of chromatin compartments is hypothesised to be driven by phase separation (5). By analysing chromatin contact maps at kilobase resolution Dixon et al. were able to identify a finer level of chromatin organisation known as Topologically Associating Domains (TADs). TADs are regions of the genome characterised by a high degree of self-interaction within the length of the TAD, and a low degree of interactions with regions outside of the TAD even if they are a similar distance away (6). TAD boundaries are enriched for CCCTC-binding factor (CTCF) binding sites and are thought to be formed by active loop extrusion, during which DNA is extruded through cohesion (forming a loop) until the extrusion is stalled by a pair convergently orientated CTCF binding sites at the TAD boundaries (5–8). Comparisons of TADs between tissues suggested that they are largely tissue invariant (4,6,9). It has also been proposed that a third category of chromatin organisation exists, which nests within TADs, these ‘sub-TADs’, are formed by the same mechanisms as TADs but have weaker insulation and are more likely to vary depending on the cell type (10). However, it is currently unclear whether sub-TADs constitute functionally different structures to TADs (10).

It has been proposed that TADs constitute functional units in the genome, important for correct regulation of gene expression. TADs co-localise regulatory elements with their target genes, and are thought to promote co-regulation of the genes they contain, creating “gene regulatory domains” (11). By inserting regulatory sensors along the length of the genome Symmons et al. found evidence that the activity of enhancers is split into regulatory domains which highly correlate with TADs (12). This provided experimental evidence that TADs potentially facilitate enhancers to carry out “non-specific” co-regulation of all genes in the TAD (11, 12). Simultaneously, TADs are thought to insulate genes from aberrant regulation by regulatory elements outside the TAD (enhancer hijacking) (11). Several examples of congenital disease have been linked to TAD boundary disruptions allowing enhancer hijacking demonstrating that at least in these cases, TAD boundaries are essential for proper gene regulation (13, 14). TAD boundaries are also able to block the spread of transcription, and repressive chromatin (11). It has been observed that the unit of compartment switching in the genome tends to be a single or series of TADs (15). Adding to this picture, it has been suggested that genes within the same TAD have highly correlated expression patterns (16–19). This has led to the speculation that genes which are required for specific processes may be contained within the same TAD to allow them to share regulatory programs and efficiently switch between the active and inactive compartments (20). Studies have already indicated that some TADs may be enriched for lineage specific genes (20, 21), but the global relationship between TADs and gene function is yet to be fully understood.

It has long been known that the linear order of genes in the genome is non-random with respect to gene function. Genes that are close together in linear space are more likely to have correlated expression patterns (22), and share pathways and protein-protein interactions (PPI) (23). Genes within TADs are by definition also close together in the linear genome therefore the linear proximity between genes is an important confounding factor when studying the similarity of genes that share a TAD. It is also possible that the increased similarity of genes that are proximal in the linear order occurs on a similar scale to TADs. By promoting co-regulation of genes, TADs could explain the increased functional similarity between proximal genes.

We hypothesised that TADs form functional units and therefore genes within them are more likely to share functional annotations than can be explained by linear proximity alone. In order to test this hypothesis we utilised some of the highest quality mammalian Hi-C data currently in the public domain (24) and annotated TADs using two TAD calling algorithms; Arrowhead and TopDom (23, 24). We first assessed the relationship between TADs and gene paralogy as well as constraint. Then, focusing on TADs most likely to have been formed by loop extrusion, we assessed the functional relatedness of non-paralogous protein coding genes within them, using four functional annotations: expression correlation, Gene ontology (GO) semantic similarity, shared pathways and PPI.

## RESULTS

### Characteristics of TADs in cortical neurons and embryonic stem cells (ESCs)

We analysed ESC and cortical neuron Hi-C data from Bonev et al. (24) using the Juicer pipeline (25). This data represents some of the highest quality mammalian Hi-C data currently in the public domain. Throughout this work we have focused on autosomal TADs, so unless explicitly stated TADs refers to autosomal TADs only. We annotated 8371 (median size 0.29Mb) and 16002 (median size 0.11Mb) TADs in ESCs, and 8001 (median size 0.32Mb) and 13835 (median size 0.12Mb) TADs in cortical neurons, using two TAD callers Arrowhead (25) and TopDom (26), respectively (Figure 1A-B). We detected more, but smaller TADs with TopDom than with Arrowhead for both cell types. Our results confirm the finding from Bonev et al. (found using the directionality index TAD calling method) that there are more and smaller TADs in ESCs than cortical neurons (24).

**Figure 1:**
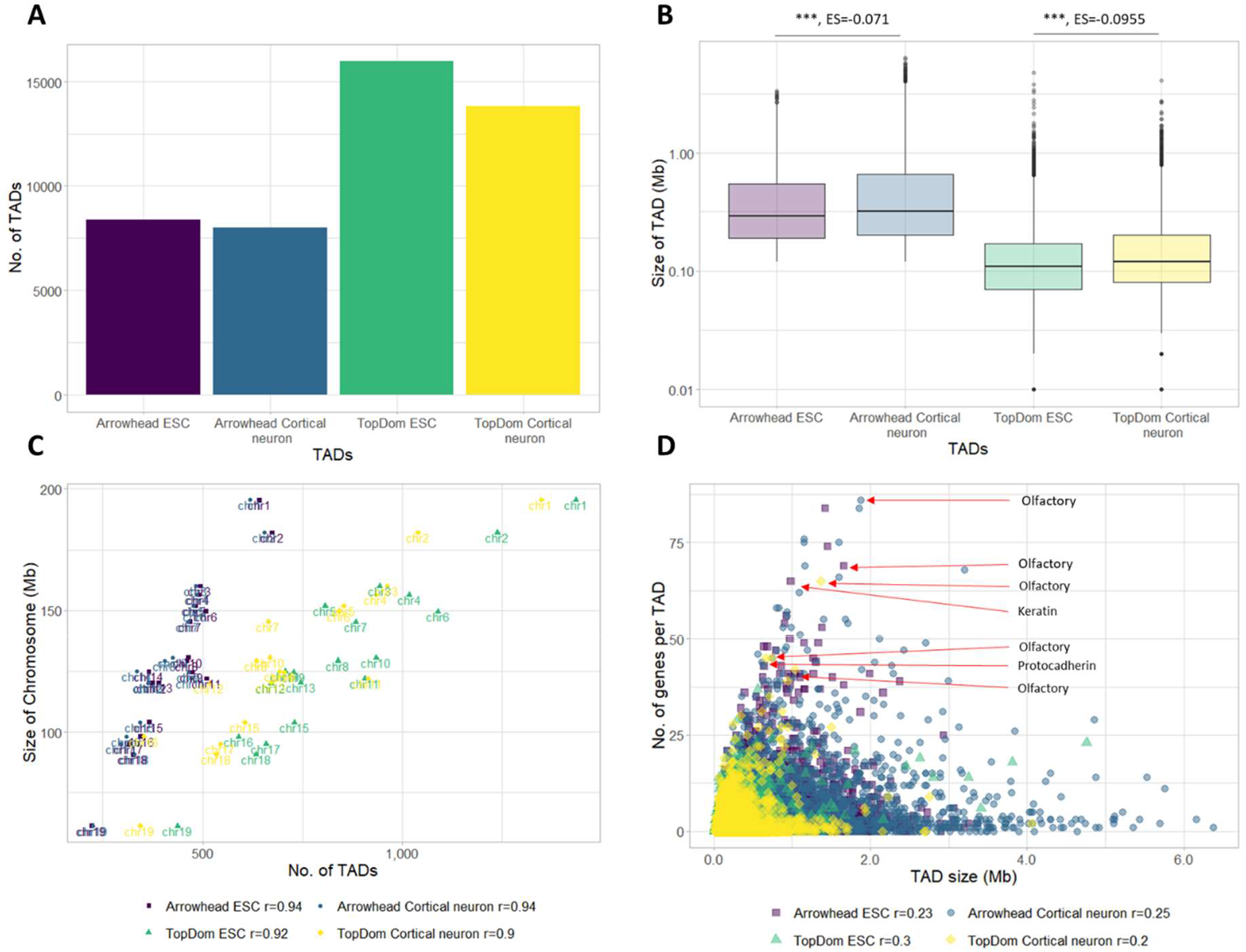
Features of autosomal TADs in ESCs and cortical neurons. A) The number of TADs called using Arrowhead and TopDom in ESCs and cortical neurons. Arrowhead calls fewer TADs than TopDom in both cell types. More TADs are called in ESCs than cortical neurons with both TAD callers. B) Size of TADs called using Arrowhead and TopDom in ESCs and cortical neurons (plotted on a log10 scale). Arrowhead calls larger TADs than TopDom in both cell types. Both Arrowhead and TopDom call significantly smaller TADs in ESCs than cortical neurons (Wilcoxon test, p-value: p<0.001 = ***, p<0.01 = **, p<0.05 = *, ES= Effect size calculated using r for Wilcoxon). C) The number of TADs per chromosome is strongly correlated with the size of the chromosome. D) In both cell types and with both TAD callers most TADs have few genes. Overall, there is a low correlation between TAD size and gene number. Several TADs containing many genes were further investigated and found to contain multiple members of large gene families (annotated).

To learn more about the distribution of TADs in the genome we looked at their association with chromosomes and protein-coding genes, matching the expected null, we observed that, for both TAD callers and datasets, the number of TADs on a chromosome correlates strongly with the size of the chromosome (Figure 1C) (r range 0.9-0.94).

Most TADs contain relatively few genes (mean number of genes: ∼2.74 and ∼3.49 (Arrowhead), and ∼0.95 and ∼1.08 (TopDom) in ESCs and cortical neurons, respectively) and there is little correlation between the number of genes within a TAD and the TAD size (r range 0.2-0.3) (Figure 1D). We investigated several TADs which contained a large number of genes and found that they contained genes from large paralog families e.g. olfactory genes and protocadherin (Figure 1D). This is consistent with previous studies which have noted that genes from these functional families tend to fall within the same TAD, likely due to their shared regulatory requirements (11).

### TAD randomisation

In order to globally assess the functional similarity between genes in the same TADs we sought to synthesise “random TADs” representing the null distribution. We developed two randomisation strategies in order to generate two null distributions controlling for different possible confounding signals. In the first randomisation strategy, which we refer to as random TADs, we maintained the basic structure of real TADs i.e. TAD size, number of genes within the TAD and the approximate TAD overlap structure. This allowed us to control for the influence of linear gene order and distance which are known to correlate with gene functional similarity (22, 23). In this randomisation strategy, the position of each TAD was randomised within the same chromosome to a new region of the same size, containing the same number of genes as the original TAD. For TopDom random TADs, TAD overlapping was prevented, reflecting the non-overlapping structure of TopDom TADs. For Arrowhead random TADs, if the new random TAD overlapped an existing random TAD this was controlled in order to favour “nested” TADs, thereby approximating the global TAD overlap structure seen in Arrowhead TADs (see Methods) (Figure 2B, Figure 3A). In the second randomisation method, which we refer to as random genome TADs, we again maintained the basic TAD structure but removed signal attributed to the linear gene. In order to do this, the positions of TADs were maintained but the order of the genes in the genome was randomly shuffled within each chromosome (Figure 2C). Using both randomisation strategies allows us to disentangle the functional similarity of genes within the same TAD from the functional similarity which can be attributed to proximity in the linear genome. Each TAD randomisation method was run 100 times for each cell type and each TAD caller.

**Figure 2:**
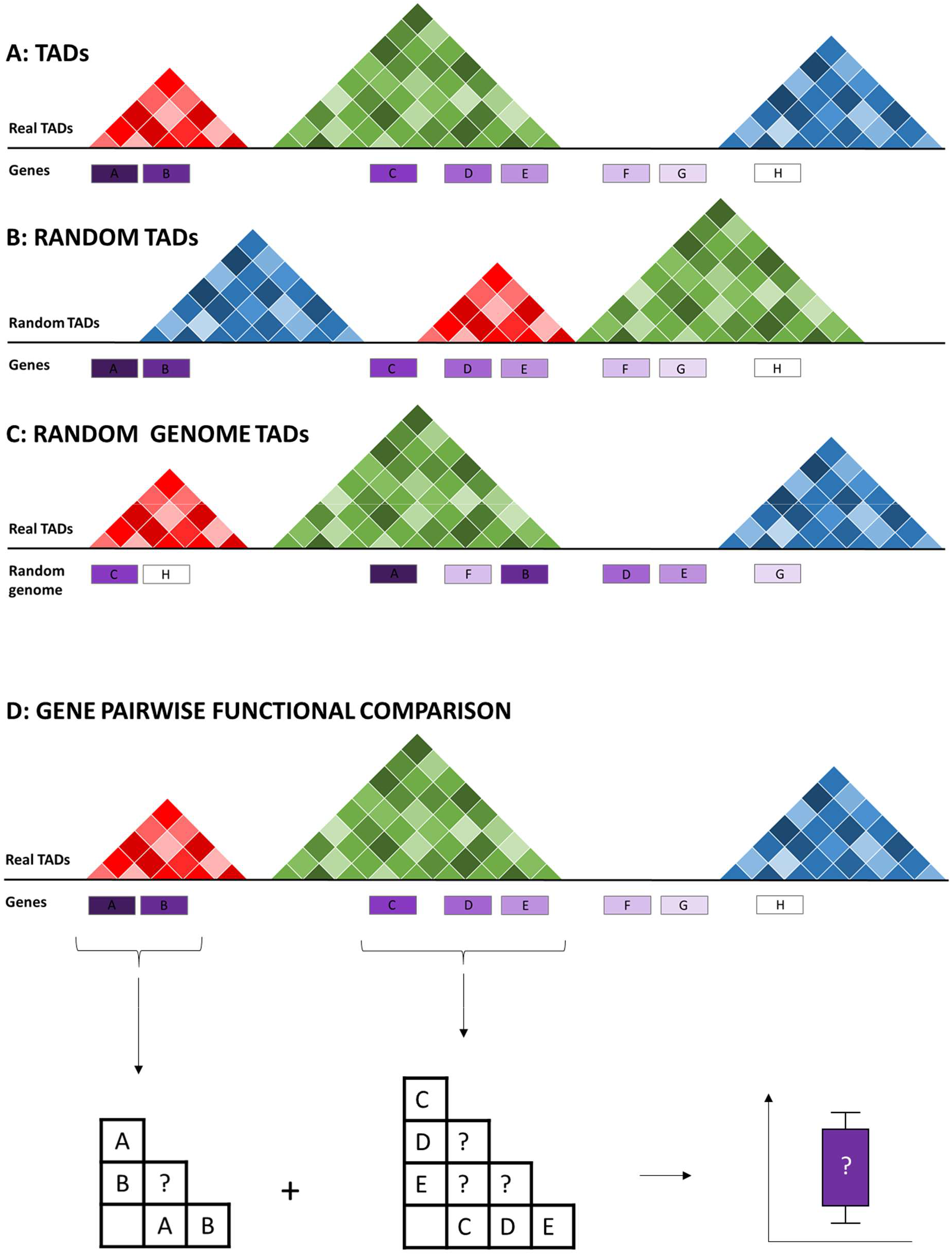
Randomization and functional analysis procedure. A: Schematic representing the structure of annotated TADs. B: Null dataset one: Random TADs. TADs were randomised within the same chromosome by selecting regions of equal size to the original TAD which also contain the same number of genes, thus controlling for the effect of the linear genome. C: Null dataset two: Random genome TADs. In order to remove the effect of the linear genome another null TAD set was generated in which the TADs remained in the same positions but the order of genes on the chromosome were randomised. C: Pairwise strategy for comparing functional similarity between genes in the same TAD. All possible pairs of genes in each TAD were compared.

**Figure 3:**
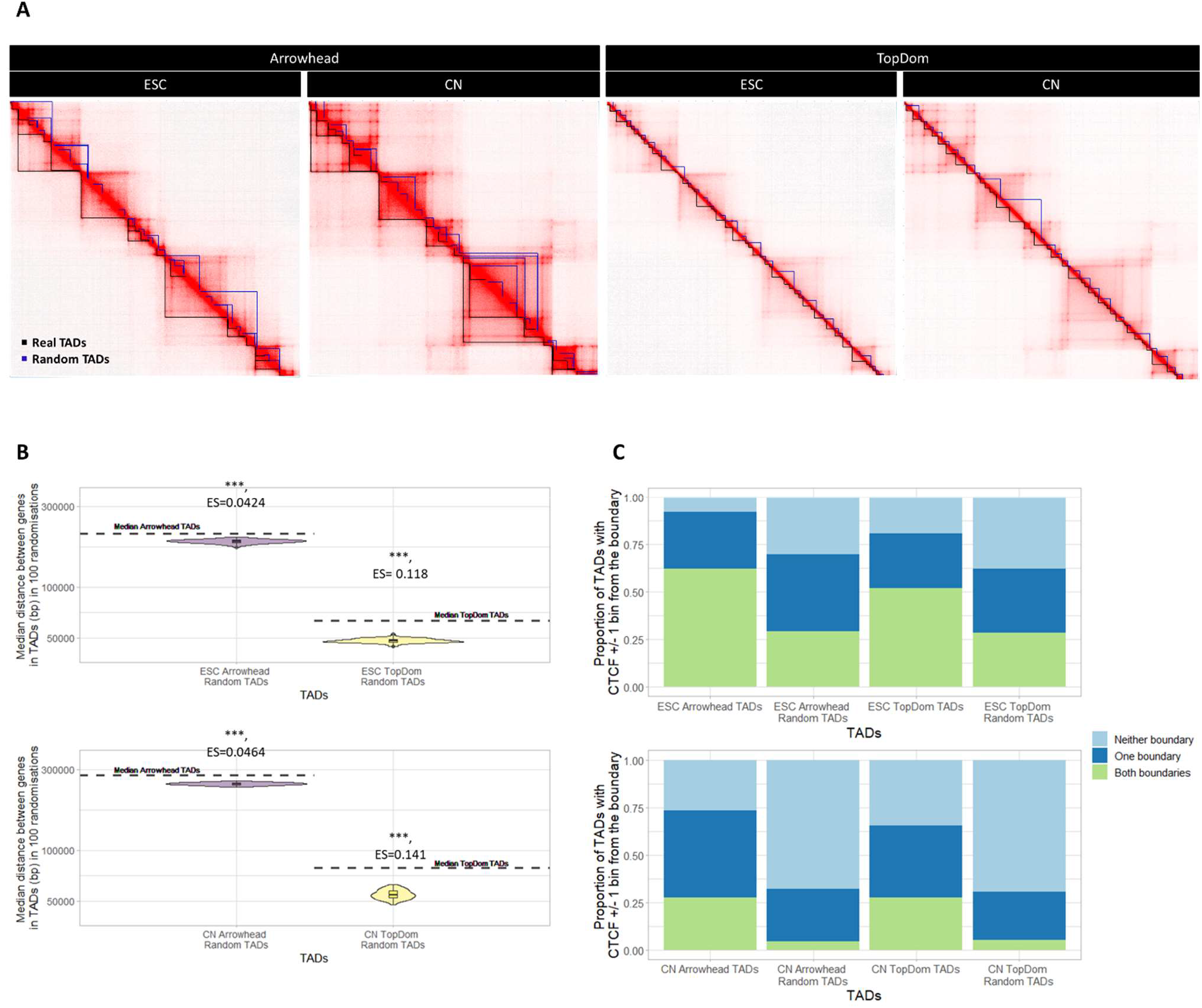
Features of autosomal TADs vs random TADs. A) TADs (black) vs an example set of random TADs (blue) shown on the Hi-C matrix for the equivalent region of Chr2 in both ESCs and cortical neurons (CN). Matrices visualised using JuiceBox. B) Median distance between genes in TADs (dotted line) vs the median distance between genes in 100 sets of random TADs (plotted on a log10 scale). Genes are significantly closer together in random TADs than TADs (Wilcoxon test, Median p-value: p<0.001 = ***, p<0.01 = **, p<0.05 = *, ES= Median effect size calculated using r for Wilcoxon). C) Proportion of TADs with a CTCF binding site within ±10kb of both boundaries, one boundary or neither boundary. As expected a greater proportion of TADs have a CTCF binding site near both boundaries than in an example set of random TADs.

In order to compare the functional similarity of genes within TADs to genes within random TADs/random genome TADs, we adopted a pairwise comparison approach (Figure 2D). For every feature investigated every possible pair of genes within a TAD/random TAD was compared in order to generate a distribution of scores. The distribution of scores for each feature in TADs was compared to each of 100 sets of random TADs and the median p-value was reported.

To assess the gene distribution within TADs, we compared the distance between genes in TADs to genes in random TADs. We found that for both TAD callers, and both cell types, genes are significantly further apart in TADs than in random TADs (Figure 3B, median p-values; Arrowhead ESC vs random TADs: p-value<0.001, Arrowhead cortical neuron vs random TADs: p-value<0.001, TopDom ESC vs random TADs: p-value<0.001, TopDom cortical neuron vs random TADs: p-value<0.001).

It has previously been shown that TAD boundaries are enriched for CTCF which is hypothesised to play a crucial role in TAD formation by loop extrusion (5–8). To assess this in our data we tested for the presence of CTCF ChIP-seq peaks near TAD boundaries vs random TAD boundaries (Figure 3C, Supplementary figure 2). We observed that ∼ 62%, 52%, 28%, 28% of ESC Arrowhead TADs, ESC TopDom TADs, cortical neuron Arrowhead TADs and cortical neuron TopDom TADs, respectively, had a CTCF ChIP-seq peak within in ±10kb of both TAD boundaries. This is compared to ∼ 29%, 29%, 4.5%, 5.3% of ESC Arrowhead random TADs, ESC TopDom random TADs, cortical neuron Arrowhead random TADs and cortical neuron TopDom random TADs, respectively. Supporting previous reports (4,6,27,28), this suggests that CTCF binding is common at the boundaries of TADs and is more prevalent than expected if TADs were randomly placed. This result also shows that more ESC TADs have a CTCF ChIP-seq peak near both boundaries than cortical neuron TADs. This could be due to a reduction in the number of chromatin domains formed by loop extrusion during differentiation. However, we noted that this still left 30%, 29%, 46%, 38% of ESC Arrowhead TADs, ESC TopDom TADs, cortical neuron Arrowhead TADs and cortical neuron TopDom TADs, respectively, which had a CTCF ChIP-seq near only one boundary and 7.9%, 19%, 27% and 35% of ESC Arrowhead TADs, ESC TopDom TADs, cortical neuron Arrowhead TADs and cortical neuron TopDom TADs, respectively, which did not have a ChIP-seq peak near either boundary. This could suggest that these domains may not be formed by loop extrusion and therefore may not conform to the popular mechanistic definition of TADs (10).

In order to assess the features of these TADs separately we split TADs into CTCF TADs (which we define as TADs with a CTCF ChIP-seq peak within ±10kb of both boundaries) and non CTCF TADs (which we define as TADs with a CTCF ChIP-seq peak within ±10kb of one, or neither boundary) (Supplementary figure 2). We compared the size of CTCF TADs and non CTCF TADs between ESCs and cortical neurons. In both CTCF TADs and non CTCF TADs we found cortical neuron TADs were significantly larger (p-value<0.001, except Arrowhead non CTCF TADs which was not significantly different) (Supplementary figure 3A-B). We also compared the distance between genes in CTCF and non CTCF TADs to random CTCF/non CTCF TADs, respectively. We found that genes are significantly further apart in both CTCF TADs and non CTCF TADs than expected in random CTCF/non CTCF TADs (p-value<0.001) (Supplementary figure 3C-D).

### TADs vs paralogy and gene constraint

We have shown examples of TADs that contain a large number of genes from the same paralogous families (Figure 1D), suggesting that genes within TADs could be more functionally similar due to shared ancestry (29). We therefore investigated whether genes within TADs are enriched for paralogous gene pairs, genome wide. To do this we assessed the proportion of paralogous gene pairs within TADs and random TADs. Similarly to Ibn-Salem et al. (30) we found a greater proportion of paralogous gene pairs fall within TADs compared to random TADs (Figure 4A, median p-value of TADs vs random TADs <0.001). This suggests that pairs of paralogous genes are more likely to fall within the same TAD than can be explained by the linear proximity of the genes alone. We further investigated this relationship and found that TADs which contain at least one pair of paralogs are more likely to be significantly larger in size than TADs with no paralogs (Figure 4B, p-value <0.001).

**Figure 4:**
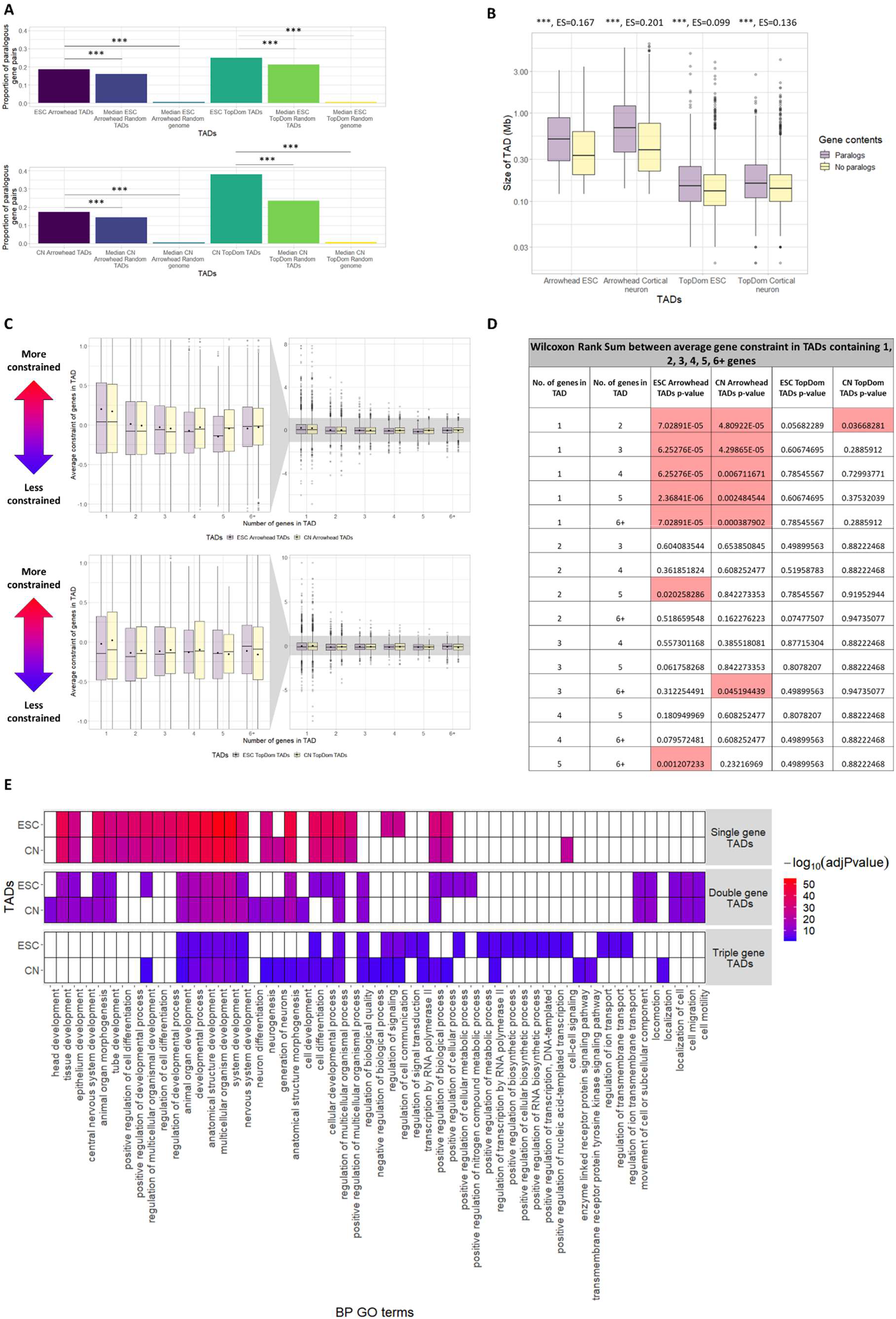
Paralogs and constraint vs autosomal TADs. A) Proportion of paralogous gene pairs in TADs, the median proportion in 100 sets of random TADs, and the median proportion in 100 sets of random genome TADs. TADs contain significantly more pairs of paralogous genes than both random TADs and random genome TADs (Fisher’s exact test, Median p-value: p<0.001 = ***, p<0.01 = **, p<0.05 = *). B) Size of TADs containing pairs of paralogs vs TADs (with >1 gene) containing no pairs of paralogs (plotted on a log10 scale). For both TAD callers and cell types, TADs which contain pairs of paralogs are larger than TADs which have no paralog pairs. (Wilcoxon test, p-value: p<0.001 = ***, p<0.01 = **, p<0.05 = *, ES= Effect size calculated using r for Wilcoxon). C) Distribution of mean constraint scores of genes occupying the same TAD. TADs are split depending on the number of genes they contain (1, 2, 3, 4, 5, 6+). Dots indicate the mean of the distribution. D) Table showing FDR corrected p-values of differences between groups in C calculated with the Wilcoxon test. Significant p-values are highlighted red. For TADs called using Arrowhead in both cell types, genes singly occupying a TAD have a significantly higher constraint score than the average constraint of genes in TADs with >1 gene. For TADs called using TopDom no significant difference is observed. E) Biological processes GO term functional enrichment of genes singly, doubly or triply occupying an Arrowhead TAD. Only the top 25 most significant GO terms passing a p-value threshold of < 0.05 (multiple testing corrected using the “gSCS” option) are shown.

When these TADs are split into CTCF TADs and non CTCF TADs we find that pairs of paralogs are significantly enriched in non CTCF TADs compared to random non CTCF TADs (median p-value of non CTCF TADs vs random non CTCF TADs <0.001). However, the opposite is true for CTCF TADs (median p-value of CTCF TADs vs random CTCF TADs <0.001 for ESC Arrowhead, ESC TopDom and cortical neuron Arrowhead and <0.01 for cortical neuron TopDom) (Supplementary figure 4). This suggests that although pairs of paralogs are enriched in TADs they are depleted in CTCF TADs, which are more likely to be “true” TADs formed by loop extrusion.

In order to further assess the impact of evolutionary forces on genes within TADs, we assessed the average constraint scores of genes in TADs. Constraint scores quantify the degree of selective constraint acting on protein coding genes, with a higher score indicating a greater strength of purifying selection (31). Selective constraint can change over evolutionary time, and we therefore considered constraint scores calculated in the mouse lineage (32). For TADs called using Arrowhead, we find that genes which singly occupy a TAD are significantly more constrained than the mean constraint of genes co-occupying TADs (Figure 4C-D). Genes singly occupying an Arrowhead TAD also have significantly higher constraint than seen in random TADs (suggesting the result cannot be explained by the structure of the linear genome alone) or random genome TADs (FDR corrected median p-value of genes singly occupying TADs vs genes singly occupying TADs in random TADs or random genome TADs <0.01, Supplementary figure 5). This suggests that genes, which singly occupy TADs, may be under higher selective constraint and more functionally important than genes which co-occupy a TAD. This might suggest that the protection from aberrant regulation of functionally important genes, implied by being in a private TAD, is under selective constraint. Interestingly, this relationship is not seen for TADs called using TopDom despite the fact that the two TAD callers detect a similar proportion of TAD containing only a single gene (∼ 21% and 20% of Arrowhead TADs, and 17% and 20% of TopDom TADs, in ESCs and cortical neurons, respectively). This could be because TopDom TADs contain a much larger percentage of TADs with no genes than Arrowhead TADs (∼ 37% and 31% of Arrowhead TADs, and 63% and 57% of TopDom TADs, in ESCs and cortical neurons, respectively). This means that there are far fewer TopDom TADs than Arrowhead TADs which contain more than one gene.

**Figure 5:**
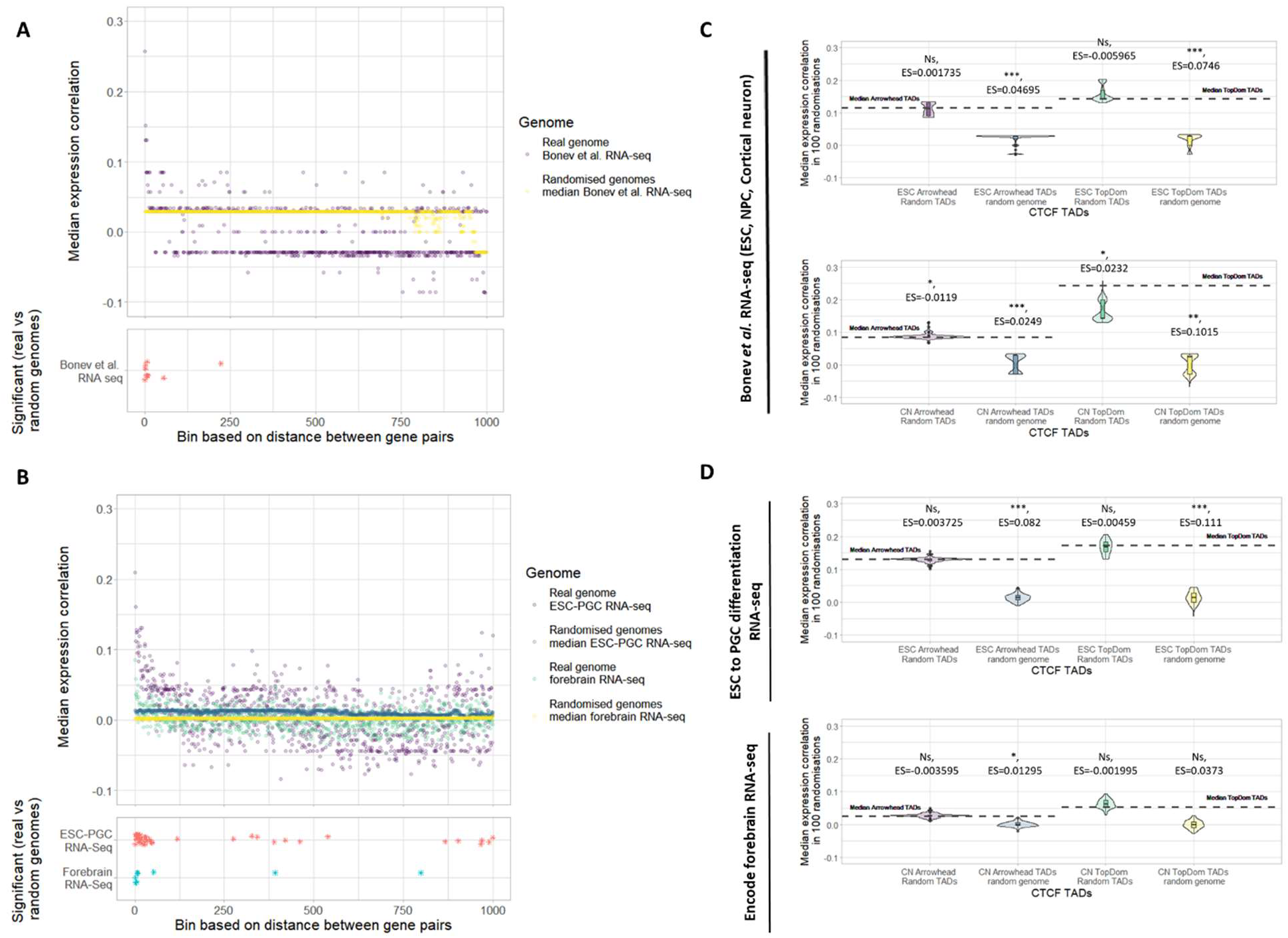
Pairwise gene co-expression in autosomal CTCF TADs. Olfactory genes and paralogous gene pairs have been excluded in all panels. A-B) Top Panel: Median expression correlation coefficient (spearman) for pairs of genes vs binned distance in the real genome and 1000 random genomes. Bottom Panel: Stars indicate bins with a significantly higher median expression correlation in the real genome vs 1000 random genomes (FDR corrected p-value<0.05). Jitter has been applied on the y axis to allow clearer visualization of close together points. A) Expression correlation coefficients were calculated using RNA-seq from two replicates each of ESC, neural progenitor cells (NPC) and cortical neuron cells. B) Expression correlation coefficients were generated using mouse ESCs differentiating to primordial germ like cells (PGC) and forebrain RNA-seq from encode. The mouse ESCs differentiating to PGC RNA-seq was generated with three replicates each of ESCs, epiblast like cells (day 2), PGC (day 4) and PGC (day 6). The forebrain RNA-seq was generated with two replicates each, of embryos of varying ages. C-D) Median expression correlation coefficient (spearman) for pairs of genes in CTCF TADs (dotted line) and median expression correlation coefficient (spearman) in 100 sets each of: random CTCF TADs and random genome CTCF TADs. CTCF TADs called using both Arrowhead and TopDom, in both ESCs and cortical neurons (CN) Hi-C (Wilcoxon test, Median p-value: p<0.001 = ***, p<0.01 = **, p<0.05 = *, NS= not significant, Median ES= Effect size calculated using r for Wilcoxon). C) Expression correlation coefficients were calculated using RNA-seq from A. D) Expression correlation coefficients were calculated using RNA-seq from B.

We next sought to test if the relationship between Arrowhead TADs and average gene constraint is observable in both CTCF TADs and non CTCF TADs. When considering only Arrowhead CTCF TADs, as seen above, we find that generally the constraint of genes in singly occupied TADs is significantly higher relative to the average constraint of genes co- occupying a CTCF TAD. On the other hand, in Arrowhead non CTCF TADs, we find a weaker relationship between genes singly occupying a non CTCF TAD and constraint (Supplementary figure 6). The difference between CTCF TADs and non CTCF TADs in terms of their enrichment for paralogous gene pairs and their relationship with gene constraint supports the possibility that TADs detected by Arrowhead and TopDom may be made up of two functional groups. As CTCF TADs are bounded by CTCF they are likely to have been formed by the process of loop extrusion and are therefore more likely to be “true TADs”. In contrast, non CTCF TADs may have been formed by other mechanisms (10).

**Figure 6:**
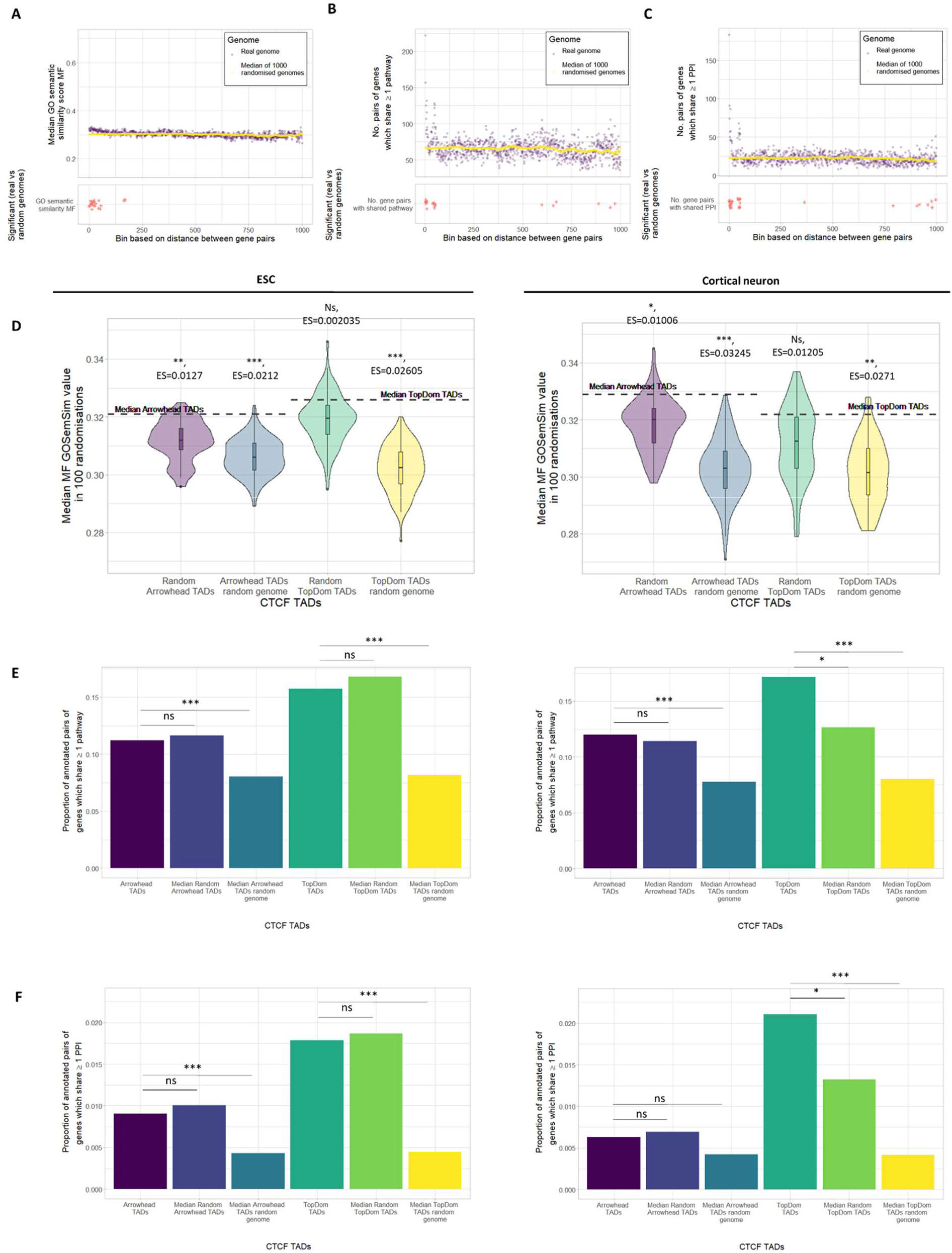
Functional similarity of pairs of genes in autosomal CTCF TADs. Olfactory genes and paralogous gene pairs have been excluded in all panels. A-C) Top Panel: Median GO semantic similarity, number of shared pathways or number of shared PPI for pairs of genes vs binned distance in the genome and 1000 random genomes. Bottom Panel: Stars indicate bins with a significantly higher functional similarity in the genome vs 1000 random genomes (FDR corrected p-value<0.05). D-F) TADs called using both Arrowhead and TopDom; in both ESC and cortical neuron Hi-C. Median P- value: p<0.001 = ***, p<0.01 = **, p<0.05 = *, NS= not significant. A) Distribution of MF GO semantic similarity for pairs of genes binned by distance in the genome vs 1000 random genomes. B) Distribution of the number of pairs of genes sharing ≥ 1 pathway binned by distance in the genome vs 1000 random genomes. C) Distribution of the number of pairs of genes sharing ≥ 1 PPI binned by distance in the genome vs 1000 random genomes. D) Median MF GO semantic similarity for pairs of genes in CTCF TADs (dotted line) compared to the distributions of median MF semantic similarity for 100 sets each of: random CTCF TADs and random genome CTCF TADs (Wilcoxon test, Median p-value, Median ES= Effect size calculated using r). E) Proportion of pairs of genes sharing ≥ 1 pathway in TADs and the median proportion of pairs of genes sharing ≥ 1 pathway in 100 sets each of: random CTCF TADs and random genome CTCF TADs (Fisher’s exact test, Median p-value. F) Proportion of pairs of genes sharing ≥ 1 PPI in TADs and the median proportion of pairs of genes sharing ≥ 1 PPI in 100 sets each of: random CTCF TADs and random genome CTCF TADs (Fisher’s exact test, Median p- value).

In order to assess which biological processes genes which singly occupy an Arrowhead TAD are involved in, we carried out a functional enrichment analysis (see Methods) using Biological process GO terms (Figure 4E). We found that genes which singly occupy an Arrowhead CTCF TAD are highly enriched for developmental processes, genes which occupy a TAD with one other gene (double occupancy) are also enriched for developmental processes but to a lesser extent and genes which occupy a TAD with two other genes (triple occupancy) are less enriched for developmental processes still. For example, “multicellular organism development” is the most significant GO term associated with singly occupied TADs in both ESCs and cortical neurons (p-value= 5.04×10^-54^ and 1.13×10^-48^, respectively), but it is less significantly associated in doubly occupied or triply occupied TADs (doubly occupied: p-value= 1.55×10^-23^ and 7.74×10^-22^, triply occupied: p-value= 1.47×10^-07^and 8.55×10^-11^ for ESC and cortical neuron, respectively). We repeated the enrichment analysis using genes which singly, doubly and triply occupy random Arrowhead TADs in order to establish whether randomly placed TADs with similar features (e.g. only one gene in the length of the TAD) have a similar pattern of enrichment (Supplementary figure 7). We found that genes that singly occupy a random TAD are also enriched for developmental processes but to a lesser degree than Arrowhead TADs. This suggests that the enrichment for developmental function observed in genes that singly occupy an Arrowhead TAD cannot be explained by the linear genome alone.

### Expression and functional similarity of non-paralogous genes in CTCF TADs

We have shown that CTCF TADs and non CTCF are unequal in their functional relevance, hence we decided to focus on the functional similarity of pairs of genes in CTCF TADs, as they are more likely to represent “true” TADs (Figure 2). Since paralogous gene pairs are highly likely to share functional similarity and we have previously assessed their relationship with TADs (Figure 4A-B, Supplementary figure 4) we excluded all pairs of paralogous genes and removed the olfactory genes (see Methods) in all functional analyses. This will allow the assessment of functional similarity between genes within TADs without recent shared ancestry.

In order to assess whether pairs of genes in the same TAD have correlated expression patterns we downloaded FPKM counts from RNA-seq expression data. RNA-seq generated during neural differentiation from Bonev et al. (24) and from the most closely matching cell types/tissues to ESCs and cortical neurons which had greater than three samples (mouse ESCs differentiating to primordial germ cell like cells (PGC) and forebrain at different embryonic stages, respectively) from Encode or GEO were used (33–36). Using these expression counts we calculated spearman’s rank correlation coefficient between pairs of genes in CTCF TADs, 100 sets of random CTCF TADs, and 100 sets of random genome CTCF TADs (Figure 5C-D). We found pairs of genes in CTCF TADs have a significantly higher expression correlation than pairs of genes in random genome CTCF TADs in 7 out of 8 comparisons (median p-value<0.05). This is an expected result because randomising the genome removes the effect of linear gene proximity. However, in 6 out of 8 comparisons we find no significant difference in expression correlation between pairs of genes in CTCF TADs and pairs of genes in random CTCF TADs (median p-value<0.05). This suggests that contrary to the majority of other studies (16–19) we find little evidence that pairs of genes sharing a TAD are more likely to have similar expression patterns than can be explained by their linear proximity. A study by Soler-Oliva et al. found that algorithmically identified co-expression domains in breast tissue/breast cancer tend not to coincide with TADs, which supports our findings (37).

Next, we sought to assess whether pairs of genes within the same CTCF TAD are more likely to share functional annotations than pairs of genes in random CTCF TADs or random genome CTCF TADs. Here, we used molecular function (MF) GO semantic similarity scores, shared pathways, and PPI (see methods). In 11 out of 12 comparisons we found that pairs of genes in CTCF TADs are significantly more similar (median p-value<0.01) in terms of functional annotation than pairs of genes in random genome CTCF TADs (Figure 6D-F).

Again, this is expected as randomising the genome removes functional similarity that can be explained by linear proximity. Using binned linear distance, we found greater similarity between pairs of genes which are in very close linear proximity than expected if genes were randomly ordered on the chromosome (Figure 6A-C). We next compared the functional annotations of genes in CTCF TADs with genes in random CTCF TADs. We found that for the majority of comparisons there was no significant difference (8 out of 12 comparisons, median p-value<0.05) (Figure 6D-F). Pairs of genes in Arrowhead ESC CTCF TADs and Arrowhead cortical neuron CTCF TADs have significantly more similar MF GO terms than pairs of genes in random CTCF TADs (median p-value <0.01 and <0.05, respectively). A similar trend in MF GO similarity was observed for all other CTCF TADs compared to random CTCF TADs but the difference wasn’t significant in the other 2 comparisons (median p-value <0.05). This could indicate that pairs of genes in CTCF TADs have slightly more similar MF GO term annotations than pairs of genes in random CTCF TADs. However, perhaps this is limited to few TADs as the increase in similarity is very small and often not significant. Pairs of genes in cortical neuron TopDom CTCF TADs are significantly more likely to share a pathway or PPI than pairs of genes in random CTCF TADs (median p-value<0.05). However, the trend in the difference in proportion of gene pairs sharing a pathway or PPI between cortical neuron Arrowhead CTCF TADs and random CTCF TADs is inconsistent and the opposite trend was observed for all ESC CTCF TADs compared to ESC random CTCF TADs. This could suggest a cell type difference in CTCF TAD structure in which pairs of genes sharing a pathway/PPI are more likely to fall within the same CTCF TAD in differentiated cells like cortical neurons compared to undifferentiated cells like ESCs. However, the inconsistency of the results suggests this effect is likely to be small and perhaps driven by few TADs. Overall, we find the biggest contribution to the functional similarity between pairs of genes in CTCF TADs can be attributed to their linear proximity in the genome. When we control for linear proximity we find a less consistent picture but in the majority of comparisons, pairs of genes in CTCF TADs are no more likely to be functionally similar than if CTCF TADs were randomly placed.

## DISCUSSION

TADs are thought to co-localise regulatory elements and their target genes, insulate genes from off-target enhancer interactions, and block the spread of genome activation (11). Due to these findings we hypothesised that genes sharing a TAD would be more likely to be co-regulated. This is because in the absence of further insulation/specificity, enhancers may be able to “scan” all regulatory elements in the TAD. We hypothesised that if this is the case one might expect genes within TADs to have higher co-expression and greater functional similarity than can be explained purely by the proximity of genes in the linear order of the genome.

Similarly to previously described work by Ibn-Salem et al. (30) we found that pairs of paralogous genes are more likely to fall within the same TAD than expected if TADs are randomly placed within chromosomes (Figure 4A). This presents a clear case in which TADs contain functionally similar genes and could reflect the need for paralogs to share regulatory elements. However, we go on to show that TADs can be split into CTCF TADs and non CTCF TADs, and whilst pairs of paralogous genes are more likely to fall within non CTCF TADs than randomly placed non CTCF TADs, paralogous gene pairs are depleted in CTCF TADs (Supplementary figure 4). This suggests that paralogs are depleted in domains formed by loop extrusion which may be more likely to represent ‘true’ TADs.

To further investigate the relationship between gene evolution and TADs, we analysed the average gene constraint of genes sharing TADs. We found that genes which singly occupy a TAD (called by Arrowhead) are statistically more constrained than genes in TADs with multiple genes. This suggests that genes in TADs on their own are less tolerant to mutation and therefore more functionally important. This is supported by Muro et al. (38) who recently found that genes which singly occupy a TAD are more likely to be associated with disease. When we separated TADs into CTCF TADs and non CTCF TADs we found that this relationship is stronger for CTCF TADs compared to non CTCF TADs (Supplementary figure 6). These results could indicate that there is a selective pressure for functionally important genes to fall privately within CTCF TADs (formed by loop extrusion) providing them with strong insulation from aberrant regulation. This selective pressure may be weaker for non CTCF TADs which may not have been formed by loop extrusion and therefore may not be as insulated. In contrast, we don’t see this relationship at all for TopDom TADs, we propose that this may be due to the smaller size of TopDom TADs which could reflect a scale more similar to that of sub-TADs (10, 39). It is also worth noting that TopDom annotates the entire genome with TADs (compared to Arrowhead, which calls them sporadically) therefore if regions exist in the genome which have no TADs, TopDom will still attempt to call them, this could increase noise in TADs called by TopDom. The differences shown here between TADs called using Arrowhead and TopDom highlight the importance of ensuring findings are robust to the choice of TAD caller.

Our results indicate that there is little evidence for an increase in expression correlation or functional annotation similarity in genes sharing a TAD. In general, we found no difference in expression correlation between pairs of non-paralogous protein coding genes in CTCF TADs vs random CTCF TADs. This is contrary to previous findings (16–19). We also found that pairs of non-paralogous protein coding genes within CTCF TADs are largely not more similar in functional annotation than in random TADs. This suggests that globally TADs are not associated with a higher degree of co-regulation between the genes they contain.

Together our results suggest TADs play a stronger role in insulating genes from aberrant regulation rather than promoting co-expression of genes within a TAD. We speculate that our results are more compatible with a model of TADs in which enhancers are prevented from interacting with all the genes within the same TAD by evolved enhancer-promoter specificity or further insulation in the form of tissue/cell type specific sub-TADs. If this is the case, disease associated variants within a TAD may have deleterious consequences by misregulating their normal target gene, but also may acquire gain-of-function regulation of other genes within the TAD.

Although referred to throughout as non CTCF TADs because they were identified by published TAD callers these domains may not represent ‘true’ TADs based on the prevalent definition (domains formed by loop extrusion). Instead, it is possible that these domains represent other domain categories e.g. compartmental domains. We therefore suggest that perhaps not all TADs called by TAD callers represent TADs. The results presented here support the assertion made by Beagan et al. (10) that it is important to separate domains formed by different mechanisms because they are likely to have different functional properties.

## CONCLUSIONS

Our results suggest a limited role for TADs in promoting co-regulation of the genes within them. We find evidence that pairs of paralogous genes fall within TADs more often than random TADs. However, we find that pairs of paralogous genes are only enriched in non CTCF TADs. The functional differences observed between CTCF and non CTCF TADs may reflect the possibility that non CTCF TADs are more similar to other types of chromatin domain (e.g. compartmental domains) than TADs (defined loop extrusion). We find little evidence that non-paralogous protein coding genes within the same CTCF TAD are more likely to have correlated expression patterns or similar functional annotations than non-paralogous protein coding genes in random TADs. This suggests that TADs formed by loop extrusion do not have a global association with co-regulation and the formation of “gene regulatory domains”. We find evidence that genes that singly occupy a CTCF TAD have significantly higher constraint. This suggests that these genes may be more functionally important and TADs formed by loop extrusion may be acting to insulate them from aberrant regulation. Overall, our results suggest a stronger role of TADs in regulatory insulation than promotion of co-regulation.

## METHODS

### Topologically associating domains

Mouse ESC and Cortical neuron Hi-C data published in (24) was downloaded from Gene Expression Omnibus (GEO) (accession number: GSE96107). These datasets represent two of the high resolution mammalian Hi-C datasets published to date. Hi-C data was analysed using the Juicer analysis pipeline aligning to the mm10 genome build (25). Parameters within Juicer were selected so that contacts with a mapping quality (MAPQ) below 30 were filtered. For each cell type all replicates were run through the Juicer pipeline separately and were combined using the “mega” option in Juicer. Hi-C data was binned at 10kb and Vanilla coverage (VC) normalisation was employed. The sex chromosomes where excluded from analysis throughout.

It has been shown that algorithmically determined TADs can vary widely depending on the TAD caller used (40–42). In order to make sure that results are robust to the choice of TAD caller, TADs were called using Arrowhead and TopDom which were both run using default parameters at 10kb. Arrowhead calls larger TADs, which can overlap whereas TopDom calls smaller non-overlapping TADs. TADs were called on Hi-C maps made from merged replicates.

It is widely suggested that TADs are formed by a loop extrusion process involving convergent CTCF bound at TAD boundaries and cohesion (10). Where indicated, TADs have been split into CTCF TAD or non CTCF TADs. In order to do this ESC and cortical neuron CTCF ChIP-seq peaks (generated alongside the Hi-C data (24)) were downloaded from GEO (GEO accession number: GSE96107). TADs where both boundaries were within ±1 bin (10kb) of a CTCF peak were considered to be “CTCF TADs”, the equivalent TADs in random TADs or random genome TADs were used for comparison. Whereas, TADs with only one boundary or neither boundary within ±1 bin (10kb) of a CTCF peak were considered to be “non CTCF TADs”.

### Overlapping TADs with protein coding genes

Ensembl IDs of mouse genes and mm10 coordinates were downloaded from BioMart (43) and non-protein coding genes were filtered out. Using bedtools intersect (44), protein coding genes were overlapped and assigned to a TAD if their start and end position fell within the same TAD. This TAD-gene mapping method is more stringent than previously used (16,17,38) but it allows us to focus on genes which can be confidently assigned to a TAD and controls for the possibility that genes which overlap a TAD boundary may have different features. This is especially important given recent evidence has shown that TAD boundaries are often not “sharp”, instead boundaries can span “zones of transition” meaning that it may not be possible to confidently assign genes spanning a TAD boundary to one TAD or the other (45)

In order to assess the functional similarity between pairs of genes in TADs, where stated olfactory genes were removed from analysis. The olfactory genes have undergone a significant expansion in the mouse vs human genome. The human genome contains ∼800 olfactory genes (of which <400 are functional), whereas the mouse genome contains ∼1400 olfactory genes (of which <1050 are functional) (46). Therefore, in order to make the findings of this study more relevant to human biology the olfactory genes were masked. To achieve this MGI IDs associated with Olfactory genes were downloaded by identifying any gene associated with the GO term: “olfactory receptor activity” (47–49) and IDs were converted to ensembl IDs using BioMart, ensembl IDs were used to remove genes from the analysis (1133 genes in total) (43).

### TAD randomisation and Genome randomisation

We synthesised “random TADs” to serve as a null distributions to compare with TADs. To do this we generated two TAD randomisation strategies, each controlling for a different possible confounding signals. In the first strategy, which we refer to as “random TADs”, the position of each TAD was randomised within the same chromosome so that each TAD was randomly assigned to a new region of the same size as the original TAD. The new region was accepted only if it contained the same number of genes as the original TAD. TopDom TADs are non-overlapping, so overlapping was prohibited in random TopDom TADs. For each TAD, if a new region satisfying the criteria could not be found after 10,000 attempts, the TAD was excluded from the random TAD set. For TADs called using Arrowhead, we observed that the overlap structure was far more likely to be “nested” (one TAD falls completely within another) than “non-nested” (TADs overlap incompletely, with only part of the TAD falling within the bounds of the other) (Supplementary figure 1). To approximate this overlap structure in random TADs, every proposed new random TAD position was checked to see if it overlapped any existing random TADs. If it overlapped an existing random TAD in a “nested” fashion the overlap was always permitted. However, if it overlapped an existing random TAD in a “non-nested” fashion, the new position was accepted with 10% probability, thereby minimising this type of overlap. As with the random TopDom TADs, if a position fulfilling this criteria could not be found after 10,000 attempts, the TAD was excluded from the random TAD set. In the second randomisation method, which we refer to as “random genome TADs” the position of TADs was maintained along with the number of protein coding genes within them, but the order of the protein coding genes on each chromosome was randomised. TAD randomisation methods were implemented using pybedtools (50).

TADs called using Arrowhead and TopDom were randomised 100 times each, generating 100 sets each of: random Arrowhead TADs, random TopDom TADs, Arrowhead random genome TADs and TopDom random genome TADs. Since, during the generation of each random TAD set, the algorithm randomises the position of each TAD in turn, the order of TADs was shuffled before the generation of every random TAD set. In order to test if 100 randomisation was enough, we plotted the median value of each measure investigated in this study with each added random TAD/random genome TAD set. For most measures the median begins to converge at fewer than 100 randomisations (Supplementary figure 8).

To test how our TAD randomisation algorithm performs compared to other recently published methods we created TAD randomisation algorithms based on the descriptions in Nora et al. 2012 (16) and Rao et al. 2014 (4). We used these algorithms to create example random TAD sets and compared them to an example random TAD set created using our method (Supplementary figure 1). In brief, in the Nora et al. 2012 method, each TAD was randomised to a region on the same chromosome which contains the same number of genes and is the same length or smaller than the original TAD. We adapted this method to prevent overlapping when randomising TopDom TADs, if a non-overlapping TAD could not be placed after 10,000 attempts it was excluded from the TopDom random TAD set. In the Roa et al. 2014 method, each TAD was randomised to a new position on the same chromosome but prevented from overlapping any gaps in the mm10 assembly (mm10 gaps were downloaded from the UCSC table browser (51)). This method was adapted to prevent overlapping when randomising TopDom TADs. Again, we set a cut off of 10,000 attempts to place each TAD before it was excluded from the TopDom random TAD set.

We compared random TADs generated using our method to random TADs generated using the Nora et al. 2012 method and the Roa et al. 2014 method (Supplementary figure 1). Since the Nora el al. 2012 method does not generate any TADs containing zero genes, TADs with zero genes were removed from all TAD sets before comparison. Regardless of the randomisation method used, we observed that the distance between genes in random TADs was always significantly different to the distance between genes in TADs. However, the effect size of these differences was smallest for Arrowhead random TADs produced by our method and second smallest for TopDom random TADs produced by our method (Rao et al. 2014 produces the smallest effect size for TopDom random TADs). This suggests that for this feature, random TADs produced by our method are the closest of the three methods to real Arrowhead TADs and second closest for TopDom TADs (Supplementary figure 1A).

We next compared the number of genes in random TADs. Again, TADs with zero genes were removed before comparison. Since in our randomisation method and in the Nora et al. 2012 method, random TADs must contain the same number of genes as the original TADs, we did not observe any difference between the number of genes within TADs and random TADs using these methods. However, we did observe a significant difference (p-value<0.001) between the number of genes in random TADs generated by the Rao et al. 2014 method and TADs (except for TopDom ESC TADs which were not significantly different) (Supplementary figure 1B). Therefore, random TADs produced using our method and the Nora et al. 2012 method best reflect the number of genes in TADs.

The overlap structure of real Arrowhead TADs favours nested TADs. To assess how well each randomisation method approximates the overlap structure of Arrowhead TADs, we selected all TADs/random TADs which were involved in any type of overlap. We then annotated them according to whether they were involved in nesting overlaps, non-nesting overlaps or both. We found that our randomisation method best approximates the overlap structure of Arrowhead TADs. Random Arrowhead TADs generated using the Rao et al. 2014 or the Nora et al. 2012 method contain more non-nesting overlaps than Arrowhead TADs (Supplementary figure 1C).

### Pairs of genes with shared ancestry

The functional similarity of genes within a TAD was measured by assessing the similarity of every possible pair of genes within the same TAD. Gene pairs which have shared ancestry i.e. paralogs, are expected to be very functionally similar. In order to assess whether genes within TADs are more functional similar irrespective of shared ancestry, (where stated) paralogous gene pairs were removed from the analysis. To do this mouse paralogous gene pairs were downloaded from BioMart (Ensemble release 98) (43).

### Constraint score

Mouse gene constraint was assessed as a nonsynonymous z-score (32), calculated between 36 strains of mice commonly used for genetic research (52). In brief, constraint was quantified for each gene as the deviation of the observed number of nonsynonymous variants relative to the expected number given no selection, which was determined by the average rate of synonymous fixation in the population sample. Genes that have a greater relative depletion of nonsynonymous variants are considered more constrained by negative selection.

### Functional enrichment analysis

Functional enrichment analysis was undertaken using the g:profiler r package (version e100_eg47_p14_021df73) (53) and Biological processes GO terms. The plots include the (max) top 25 most significant GO terms passing a p-value threshold of < 0.05 (multiple testing corrected using the “gSCS” option). The similarity between each of the significant GO terms was calculated using the r package GOSemSim (54) (using the “Jiang” similarity method as the “measure” parameter and “NULL” as the “combine” parameter). These similarity scores were then used as input into hclust clustering, the order resulting from clustering was used to order the GO terms in the plots.

### Gene co-expression

FPKM counts from RNA-seq data generated alongside the Hi-C data was downloaded from GEO (GEO accession number: GSE96107). The data consisted of two replicates each for ESC, neural progenitor cells (NPC), and cortical neurons. Genes with an FPKM value <1 were treated as having 0 expression. Gene co-expression was calculated across all 6 samples using spearman’s rank correlation coefficient between all pairs of genes. Correlation coefficients calculated from this data indicate the similarity of expression over three cell types.

To guard against the possibility that TAD structure may differ between cell types we also downloaded RNA seq data from the closest matching tissues/cell types to ESCs and cortical neurons which had at least 3 samples (required for the correlation analyses) from the Encode project (33) and GEO (35). To assess the expression correlation within cortical neuron TADs we downloaded encode forebrain RNA-seq. The forebrain RNA-seq was generated with two replicates each, of embryos of varying ages. Accession numbers: ENCFF302TQO, ENCFF976OLT, ENCFF895JXR, ENCFF227HKF, ENCFF340XFQ, ENCFF484AOO, ENCFF601JPN, ENCFF413BXV, ENCFF465SNB, ENCFF567AFL, ENCFF590FAC, ENCFF745ZJF, ENCFF763GXJ, ENCFF804FTJ, ENCFF816CVP and ENCFF918QNL (33, 34). Genes with an FPKM value <1 were treated as having 0 expression. To assess the expression correlation within ESC TADs we downloaded mouse ESCs differentiating to PGC RNA-seq data. The ESCs differentiating to PGC RNA-seq data was generated with three replicates each of ESC, epiblast like cells (day 2), PGC (day 4) and PGC (day 6) (36). Processed RNA-seq data was downloaded from GEO accession GSE86903, data was provided as log2(RPM) which we converted to RPKM for analysis. Genes with an RPKM value <1 were treated as having 0 expression.

### GO Semantic similarity

The R package GOSemSim (54) was used to calculate GO semantic similarity scores. For each pair of genes the GO terms assigned to them were compared using the Jiang method. If genes were associated with multiple GO terms, scores were combined using “best match average”. Pairs of genes where one or both have no annotated GO terms were excluded from the analysis as no score could be generated. We first calculated the similarity score between all pairs of genes in the genome using each of the MF, Biological process (BP) or Cellular component (CC) ontologies. We then plotted these scores for all autosomal genes, autosomal genes minus olfactory genes, and autosomal genes minus olfactory genes and paralogous pairs, against genomic distance in the real genome compared to the median distance in 1000 random genomes. We found that, for scores calculated with BP and CC, once olfactory genes and paralogous gene pairs have been removed there is no association between GO similarity and distance. This suggests that similarity in these scores is driven by paralogous pairs and the olfactory genes. We therefore moved forward using only MF in our analysis (Supplementary figure 9).

### Shared pathways

Kegg pathways were downloaded from org.Mm.eg.db (55). The Kegg pathway data is very sparse and many genes do not have a pathway annotation. In order to account for this, the amount of pairs of genes sharing at least one pathway annotation was considered as a proportion of all pairs of genes with at least one pathway annotation each.

### Shared protein-protein interactions (PPIs)

PPI data was downloaded for mm10 from string v11 (56). Only interactions with the mode “binding” were selected so that only direct/physical interactions (rather than functional interactions which may not require physical contact) were included. Similarly to pathways, PPI data is very sparse and many genes are unannotated. Therefore, as with pathways the amount of pairs of genes with a PPI was considered as a proportion of all pairs of genes with at least one PPI annotation each.

### Random permutation testing

For expression correlation, GO semantic similarity, proportion of gene pairs sharing a pathway and proportion of genes pairs sharing a PPI we have plotted the functional score against binned genomic distance in the real genome and compared it to the median value of the functional score in 1000 random genomes. In these analyses for each bin, we have established whether there is a significant difference between the functional score in the real genome compared to the distribution of scores in 1000 random genomes using permutation testing. For each bin this has been calculated as follows: sum(values in the random genome ≥ value in the real genome)/1000. P-values were then FDR corrected.

### Effect size

Effect size, r, was calculated using the r package rcompanion. A positive effect size indicates that the value associated with TADs is greater (than random TADs/random genome TADs) whereas a negative effect size indicates that the value associated with TADs is lesser (than random TADs/random genome TADs). The larger the value the larger the effect size.

## LIST OF ABBREVIATIONS

CN: Cortical neurons
ESC Embryonic stem cells
FPKM: Fragments Per Kilobase of transcript per Million mapped reads
GEO: Gene Expression Omnibus
GO: Gene ontology
MAPQ: MAPping Quality
MF: Molecular function
NPC: Neural progenitor cells
NS: Not significant
PGC: Primordial germ cell
PPI: Protein protein interactions
RPKM: Reads Per Kilobase of transcript per Million mapped reads
TAD: Topologically associating domains

## AVAILABILITY OF DATA AND MATERIALS

The processed datasets analysed in the current study will be made available upon publication.

## DECLARATIONS

### Ethics approval and consent to participate

Not applicable.

### Consent for publication

Not applicable.

### Authors’ contributions

HSL and MMS conceived and directed the research. HSL carried out the majority of the analysis and wrote the manuscript. SG provided statistical insight. GP calculated the constraint scores. AMM, CML and MMS provided critical feedback and supervised this work. All authors approved the manuscript.

## Acknowledgements

We would like to acknowledge the helpful feedback and comments given by Dr Christoffer Nellåker and Dr Saskia Reibe during the preparation of this manuscript.

## REFERENCES

1. Rowley MJ, Corces VG. Organizational principles of 3D genome architecture. Nat Rev Genet [Internet]. 2018 Dec 26 [cited 2019 Oct 28];19(12):789–800. Available from: http://www.nature.com/articles/s41576-018-0060-8

2. Dekker J, Rippe K, Dekker M, Kleckner N. Capturing chromosome conformation. Science [Internet]. 2002 Feb 15 [cited 2018 Nov 23];295(5558):1306–11. Available from: http://www.ncbi.nlm.nih.gov/pubmed/11847345

3. Lieberman-Aiden E, van Berkum NL, Williams L, Imakaev M, Ragoczy T, Telling A, et al. Comprehensive mapping of long-range interactions reveals folding principles of the human genome. Science [Internet]. 2009 Oct 9 [cited 2018 Feb 28];326(5950):289–93. Available from: http://www.ncbi.nlm.nih.gov/pubmed/19815776

4. Rao SSP, Huntley MH, Durand NC, Stamenova EK, Bochkov ID, Robinson JT, et al. A 3D map of the human genome at kilobase resolution reveals principles of chromatin looping. Cell [Internet]. 2014 Dec 18 [cited 2018 Aug 13];159(7):1665–80. Available from: http://www.ncbi.nlm.nih.gov/pubmed/25497547

5. Hildebrand EM, Dekker J. Mechanisms and Functions of Chromosome Compartmentalization. Vol. 45, Trends in Biochemical Sciences. Elsevier Ltd; 2020. p. 385–96.

6. Dixon JR, Selvaraj S, Yue F, Kim A, Li Y, Shen Y, et al. Topological domains in mammalian genomes identified by analysis of chromatin interactions. Nature [Internet]. 2012 Apr 11 [cited 2018 Feb 5];485(7398):376–80. Available from: http://www.nature.com/doifinder/10.1038/nature11082

7. Nuebler J, Fudenberg G, Imakaev M, Abdennur N, Mirny LA. Chromatin organization by an interplay of loop extrusion and compartmental segregation. Proc Natl Acad Sci U S A. 2018 Jul 17;115(29):E6697–706.

8. Mirny LA, Imakaev M, Abdennur N. Two major mechanisms of chromosome organization. Curr Opin Cell Biol [Internet]. 2019 Jun 19 [cited 2019 Jul 8];58:142–52. Available from: http://www.ncbi.nlm.nih.gov/pubmed/31228682

9. Schmitt AD, Hu M, Jung I, Xu Z, Qiu Y, Tan CL, et al. A Compendium of Chromatin Contact Maps Reveals Spatially Active Regions in the Human Genome. Cell Rep [Internet]. 2016 Nov 15 [cited 2018 May 16];17(8):2042–59. Available from: https://www.sciencedirect.com/science/article/pii/S2211124716314814?via%3Dihub#mmc1

10. Beagan JA, Phillips-Cremins JE. On the existence and functionality of topologically associating domains. Nat Genet [Internet]. 2020 Jan 10 [cited 2020 Jan 13]; Available from: http://www.nature.com/articles/s41588-019-0561-1

11. Dixon JR, Gorkin DU, Ren B. Chromatin Domains: The Unit of Chromosome Organization. Mol Cell [Internet]. 2016 [cited 2018 Apr 16];62:668–80. Available from: http://dx.doi.org/10.1016/j.molcel.2016.05.018

12. Symmons O, Uslu VV, Tsujimura T, Ruf S, Nassari S, Schwarzer W, et al. Functional and topological characteristics of mammalian regulatory domains. Genome Res [Internet]. 2014 Mar [cited 2018 Feb 5];24(3):390–400. Available from: http://www.ncbi.nlm.nih.gov/pubmed/24398455

13. Franke M, Ibrahim DM, Andrey G, Schwarzer W, Heinrich V, Schöpflin R, et al. Formation of new chromatin domains determines pathogenicity of genomic duplications. Nature [Internet]. 2016 Oct 5 [cited 2018 Mar 7];538(7624):265–9. Available from: http://www.nature.com/articles/nature19800

14. Scacheri CA, Scacheri PC. Mutations in the noncoding genome. Curr Opin Pediatr [Internet]. 2015 Dec [cited 2018 Mar 8];27(6):659–64. Available from: http://www.ncbi.nlm.nih.gov/pubmed/26382709

15. Dixon JR, Jung I, Selvaraj S, Shen Y, Antosiewicz-Bourget JE, Lee AY, et al. Chromatin architecture reorganization during stem cell differentiation. Nature [Internet]. 2015 Feb 19 [cited 2018 Nov 22];518(7539):331–6. Available from: http://www.ncbi.nlm.nih.gov/pubmed/25693564

16. Nora EP, Lajoie BR, Schulz EG, Giorgetti L, Okamoto I, Servant N, et al. Spatial partitioning of the regulatory landscape of the X-inactivation centre. Nature [Internet]. 2012 Apr 11 [cited 2018 Nov 23];485(7398):381–5. Available from: http://www.nature.com/doifinder/10.1038/nature11049

17. Flavahan WA, Drier Y, Liau BB, Gillespie SM, Venteicher AS, Stemmer-Rachamimov AO, et al. Insulator dysfunction and oncogene activation in IDH mutant gliomas. Nature. 2016 Jan 7;529(7584):110–4.

18. Tarbier M, Mackowiak SD, Frade J, Catuara-Solarz S, Biryukova I, Gelali E, et al. Nuclear gene proximity and protein interactions shape transcript covariances in mammalian single cells. bioRxiv. 2019 Sep 16;771402.

19. Sarnataro S, Riba A, Molina N. Regulation of transcription reactivation dynamics exiting mitosis. bioRxiv. 2020 Apr 16;2020.04.15.042853.

20. Ruiz-Velasco M, Zaugg JB. Structure meets function: How chromatin organisation conveys functionality. Curr Opin Syst Biol [Internet]. 2017 Feb 1 [cited 2019 Jan 10];1:129–36. Available from: https://www.sciencedirect.com/science/article/pii/S2452310017300173?dgcid=raven_sd_recommender_email

21. Neems DS, Garza-Gongora AG, Smith ED, Kosak ST. Topologically associated domains enriched for lineage-specific genes reveal expression-dependent nuclear topologies during myogenesis. Proc Natl Acad Sci [Internet]. 2016 Mar 22 [cited 2019 Jan 10];113(12):E1691– 700. Available from: https://www.pnas.org/content/113/12/E1691.long

22. Hurst LD, Pál C, Lercher MJ. The evolutionary dynamics of eukaryotic gene order. Nat Rev Genet [Internet]. 2004 Apr [cited 2019 Aug 23];5(4):299–310. Available from: http://www.nature.com/articles/nrg1319

23. Thévenin A, Ein-Dor L, Ozery-Flato M, Shamir R. Functional gene groups are concentrated within chromosomes, among chromosomes and in the nuclear space of the human genome. Nucleic Acids Res [Internet]. 2014 Sep 2 [cited 2019 Apr 24];42(15):9854–61. Available from: http://academic.oup.com/nar/article/42/15/9854/2435256/Functional-gene-groups-are-concentrated-within

24. Bonev B, Mendelson Cohen N, Szabo Q, Fritsch L, Papadopoulos GL, Lubling Y, et al. Multiscale 3D Genome Rewiring during Mouse Neural Development. Cell [Internet]. 2017 Oct 19 [cited 2018 Dec 7];171(3):557-572.e24. Available from: http://www.ncbi.nlm.nih.gov/pubmed/29053968

25. Durand NC, Shamim MS, Machol I, Rao SSP, Huntley MH, Lander ES, et al. Juicer Provides a One-Click System for Analyzing Loop-Resolution Hi-C Experiments. Cell Syst [Internet]. 2016 Jul 27 [cited 2018 Jul 30];3(1):95–8. Available from: https://www.sciencedirect.com/science/article/pii/S2405471216302198?via%3Dihub

26. Shin H, Shi Y, Dai C, Tjong H, Gong K, Alber F, et al. TopDom: an efficient and deterministic method for identifying topological domains in genomes. Nucleic Acids Res [Internet]. 2016 Apr 20 [cited 2018 Jun 21];44(7):e70–e70. Available from: https://academic.oup.com/nar/article-lookup/doi/10.1093/nar/gkv1505

27. Sanborn AL, Rao SSP, Huang S-C, Durand NC, Huntley MH, Jewett AI, et al. Chromatin extrusion explains key features of loop and domain formation in wild-type and engineered genomes. Proc Natl Acad Sci U S A [Internet]. 2015 Nov 24 [cited 2018 Nov 28];112(47):E6456-65. Available from: http://www.ncbi.nlm.nih.gov/pubmed/26499245

28. Vietri Rudan M, Barrington C, Henderson S, Ernst C, Odom DT, Tanay A, et al. Comparative Hi- C reveals that CTCF underlies evolution of chromosomal domain architecture. Cell Rep [Internet]. 2015 Mar 3 [cited 2019 Jan 9];10(8):1297–309. Available from: http://www.ncbi.nlm.nih.gov/pubmed/25732821

29. Stamboulian M, Guerrero RF, Hahn MW, Radivojac P. The ortholog conjecture revisited: The value of orthologs and paralogs in function prediction. Bioinformatics [Internet]. 2020 [cited 2021 Apr 13];36(Suppl 1):I219–26. Available from: /pmc/articles/PMC7355290/

30. Ibn-Salem J, Muro EM, Andrade-Navarro MA. Co-regulation of paralog genes in the three- dimensional chromatin architecture. Nucleic Acids Res. 2017;45(1):81–91.

31. Bartha I, Di Iulio J, Venter JC, Telenti A. Human gene essentiality. Nat Rev Genet [Internet]. 2018 Oct 30 [cited 2020 Sep 24];19(1):51–62. Available from: www.nature.com/nrg

32. Powell G, Simon M, Pulit S, Mallon A-M, Lindgren C. Tolerance of nonsynonymous variation is closely correlated between human and mouse orthologues. bioRxiv [Internet]. 2019 Jun 3 [cited 2020 Sep 24];657981. Available from: https://doi.org/10.1101/657981

33. Davis CA, Hitz BC, Sloan CA, Chan ET, Davidson JM, Gabdank I, et al. The Encyclopedia of DNA elements (ENCODE): Data portal update. Nucleic Acids Res [Internet]. 2018 Jan 1 [cited 2020 Sep 25];46(D1):D794–801. Available from: https://pubmed.ncbi.nlm.nih.gov/29126249/

34. He P, Williams BA, Trout D, Marinov GK, Amrhein H, Berghella L, et al. The changing mouse embryo transcriptome at whole tissue and single-cell resolution. Nature. 2020 Jul 30;583(7818):760–7.

35. Edgar R, Domrachev M, Lash AE. Gene Expression Omnibus: NCBI gene expression and hybridization array data repository. Nucleic Acids Res [Internet]. 2002 Jan 1 [cited 2018 Nov 19];30(1):207–10. Available from: http://www.ncbi.nlm.nih.gov/pubmed/11752295

36. von Meyenn F, Berrens R V., Andrews S, Santos F, Collier AJ, Krueger F, et al. Comparative Principles of DNA Methylation Reprogramming during Human and Mouse In Vitro Primordial Germ Cell Specification. Dev Cell [Internet]. 2016 Oct 10 [cited 2021 Apr 7];39(1):104–15. Available from: https://pubmed.ncbi.nlm.nih.gov/27728778/

37. Soler-Oliva ME, Guerrero-Martínez JA, Bachetti V, Reyes JC. Analysis of the relationship between coexpression domains and chromatin 3D organization. PLoS Comput Biol. 2017 Sep 1;13(9).

38. Muro EM, Ibn-Salem J, Andrade-Navarro MA. The distributions of protein coding genes within chromatin domains in relation to human disease. Epigenetics and Chromatin [Internet]. 2019 Dec 5 [cited 2020 Mar 10];12(1):72. Available from: https://epigeneticsandchromatin.biomedcentral.com/articles/10.1186/s13072-019-0317-2

39. Phillips-Cremins JE, Sauria MEG, Sanyal A, Gerasimova TI, Lajoie BR, Bell JSK, et al. Architectural Protein Subclasses Shape 3D Organization of Genomes during Lineage Commitment. Cell [Internet]. 2013 Jun 6 [cited 2018 Feb 5];153(6):1281–95. Available from: http://www.ncbi.nlm.nih.gov/pubmed/23706625

40. Dali R, Blanchette M. A critical assessment of topologically associating domain prediction tools. Nucleic Acids Res [Internet]. 2017 Apr 7 [cited 2018 May 4];45(6):2994–3005. Available from: https://academic.oup.com/nar/article-lookup/doi/10.1093/nar/gkx145

41. Forcato M, Nicoletti C, Pal K, Livi CM, Ferrari F, Bicciato S. Comparison of computational methods for Hi-C data analysis. Nat Methods. 2017;14(7).

42. Zufferey M, Tavernari D, Oricchio E, Ciriello G. Comparison of computational methods for the identification of topologically associating domains. Genome Biol [Internet]. 2018 Dec 10 [cited 2018 Dec 12];19(1):217. Available from: https://genomebiology.biomedcentral.com/articles/10.1186/s13059-018-1596-9

43. Smedley D, Haider S, Ballester B, Holland R, London D, Thorisson G, et al. BioMart - Biological queries made easy. BMC Genomics [Internet]. 2009 Jan 14 [cited 2020 Mar 10];10(1):22. Available from: http://bmcgenomics.biomedcentral.com/articles/10.1186/1471-2164-10-22

44. Quinlan AR, Hall IM. BEDTools: A flexible suite of utilities for comparing genomic features. Bioinformatics [Internet]. 2010 Jan 28 [cited 2020 Sep 25];26(6):841–2. Available from: /pmc/articles/PMC2832824/?report=abstract

45. Chang LH, Ghosh S, Noordermeer D. TADs and Their Borders: Free Movement or Building a Wall? Vol. 432, Journal of Molecular Biology. Academic Press; 2020. p. 643–52.

46. Niimura Y. Evolutionary dynamics of olfactory receptor genes in chordates: interaction between environments and genomic contents. Vol. 4, Human genomics. BioMed Central; 2009. p. 107–18.

47. Carbon S, Ireland A, Mungall CJ, Shu S, Marshall B, Lewis S, et al. AmiGO: online access to ontology and annotation data. Bioinforma Appl NOTE [Internet]. 2009 [cited 2020 Mar 10];25(2):288–9. Available from: http://www.geneontology.org

48. The Gene Ontology Consortium. The Gene Ontology Resource: 20 years and still GOing strong. Nucleic Acids Res [Internet]. 2018 Nov 5;47(D1):D330–8. Available from: https://doi.org/10.1093/nar/gky1055

49. Ashburner M, Ball CA, Blake JA, Botstein D, Butler H, Cherry JM, et al. Gene ontology: Tool for the unification of biology. Vol. 25, Nature Genetics. NIH Public Access; 2000. p. 25–9.

50. Dale RK, Pedersen BS, Quinlan AR. Pybedtools: A flexible Python library for manipulating genomic datasets and annotations. Bioinformatics [Internet]. 2011 Dec [cited 2020 Sep 25];27(24):3423–4. Available from: https://pubmed.ncbi.nlm.nih.gov/21949271/

51. Karolchik D, Hinricks AS, Furey TS, Roskin KM, Sugnet CW, Haussler D, et al. The UCSC table browser data retrieval tool. Nucleic Acids Res [Internet]. 2004 Jan 1 [cited 2020 Sep 25];32(DATABASE ISS.). Available from: https://pubmed.ncbi.nlm.nih.gov/14681465/

52. Keane TM, Goodstadt L, Danecek P, White MA, Wong K, Yalcin B, et al. Mouse genomic variation and its effect on phenotypes and gene regulation. Nature [Internet]. 2011 Sep 15 [cited 2018 Dec 11];477(7364):289–94. Available from: http://www.nature.com/articles/nature10413

53. Raudvere U, Kolberg L, Kuzmin I, Arak T, Adler P, Peterson H, et al. g:Profiler: a web server for functional enrichment analysis and conversions of gene lists (2019 update). Nucleic Acids Res [Internet]. 2019 May 8;47(W1):W191–8. Available from: https://doi.org/10.1093/nar/gkz369

54. Yu G, Li F, Qin Y, Bo X, Wu Y, Wang S. GOSemSim: an R package for measuring semantic similarity among GO terms and gene products. Bioinformatics [Internet]. 2010 Apr 1 [cited 2019 Jul 17];26(7):976–8. Available from: https://academic.oup.com/bioinformatics/article-lookup/doi/10.1093/bioinformatics/btq064

55. Carlson M. org.Mm.eg.db: Genome wide annotation for Mouse. R package version 3.5.0. Bioconductor. 2017.

56. Szklarczyk D, Gable AL, Lyon D, Junge A, Wyder S, Huerta-Cepas J, et al. STRING v11: Protein- protein association networks with increased coverage, supporting functional discovery in genome-wide experimental datasets. Nucleic Acids Res. 2019 Jan 8;47(D1):D607–13.

