## Supplementary figures for "Making sense of the linear genome, gene function and TADs"

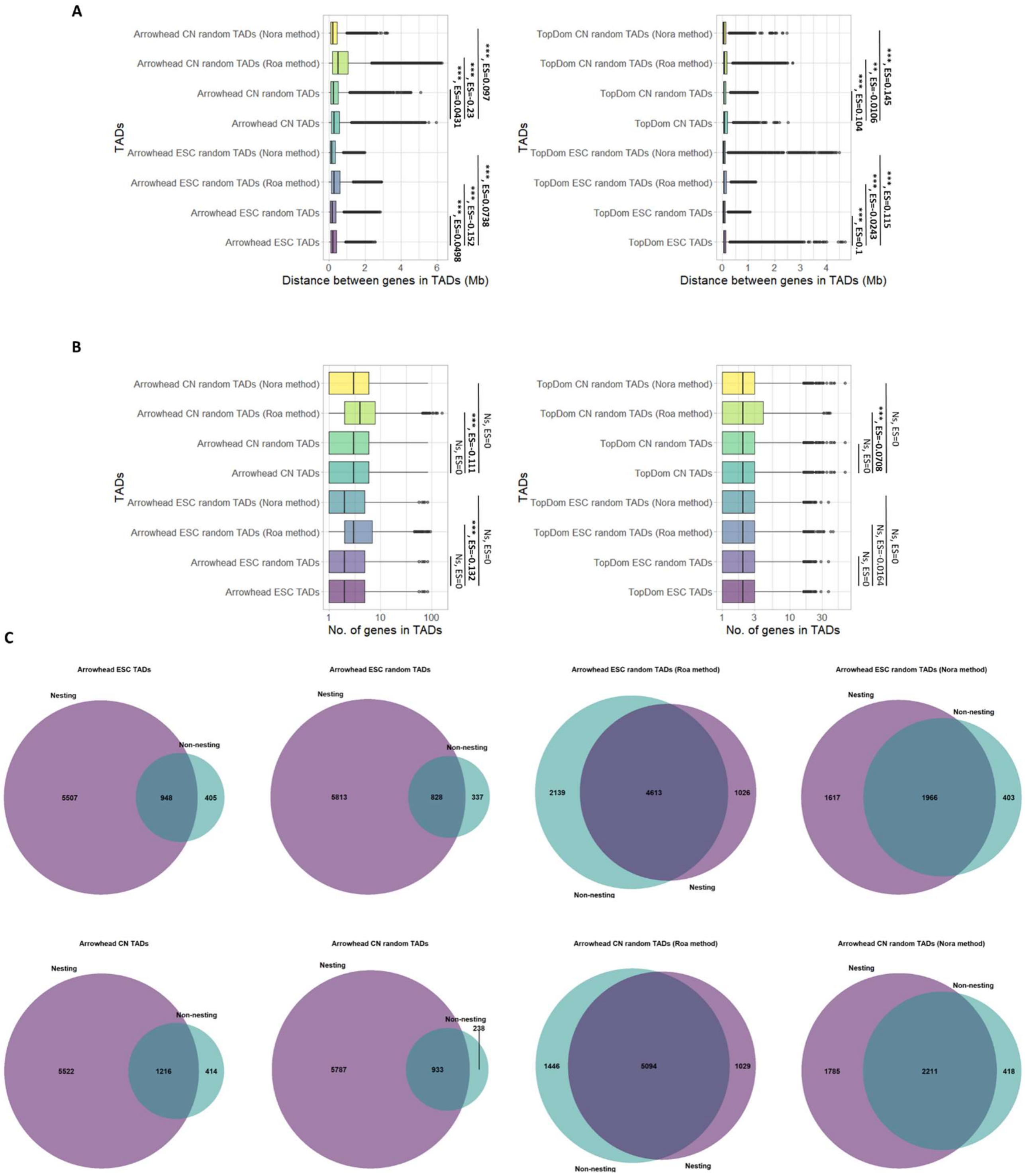


***Supplementary figure 1: TAD randomisation method compared to recently published methods (autosomal TADs).*** *Comparison of Arrowhead or TopDom TADs, and an example set of random Arrowhead or TopDom TADs generated either: using our TAD randomisation method, as described in*

*Nora et al. 2012, or as described in Roa et al. 2014. A) Distance between pairs of genes in TADs/random TADs. TADs containing no genes were excluded. The distribution of distances between pairs of genes in random TADs is significantly different from TADs, regardless of the randomisation method used (Wilcoxon test, p-value: p<0.001 = ***, p<0.01 = **, p<0.05 = *, ES= Effect size calculated using r for Wilcoxon). B) Number of genes in TADs/random TADs. TADs containing no genes were excluded. There is no difference between the number of genes in random TADs generated using either our method or Nora et al. 2012, compared to TADs. However, we find a significantly greater number of genes falling within random TADs generated as described in Roa et al. 2014 compared to TADs (except TopDom ESC TADs). (Wilcoxon test, p-value: p<0.001 = ***, p<0.01 = **, p<0.05 = *, NS= not significant, ES= Effect size calculated using r for Wilcoxon). C) Venn diagrams displaying the overlap structure of Arrowhead TADs and an example set of Arrowhead random TADs generated by each method. Non-Overlapping TADs were excluded. "Nesting" is defined as an overlap in which one TAD is contained entirely within another TAD, and "non-nesting" is defined as any incomplete overlap (i.e. only part of one TAD is contained within another TAD). The overlap structure of Arrowhead TADs is characterised by a large degree of nesting. The overlap structure of Arrowhead random TADs generated with our method has a similar overlap structure to real Arrowhead TADs. Whereas, Arrowhead random TADs generated as described by either Roa et al. 2014 or Nora et al. 2012 contain more non-nested overlaps.*

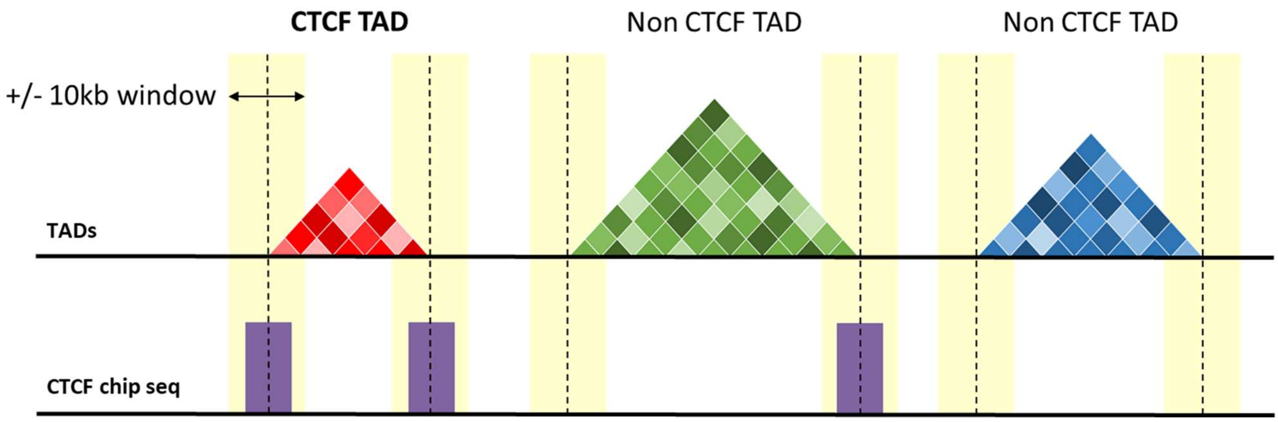


***Supplementary figure 2: CTCF TAD selection.*** *Schematic demonstrating how CTCF TADs are defined. A TAD was annotated as a "CTCF TAD" if both boundaries fall within ±10kb of a CTCF peak in CTCF ChIP-seq data from the same cell type.*

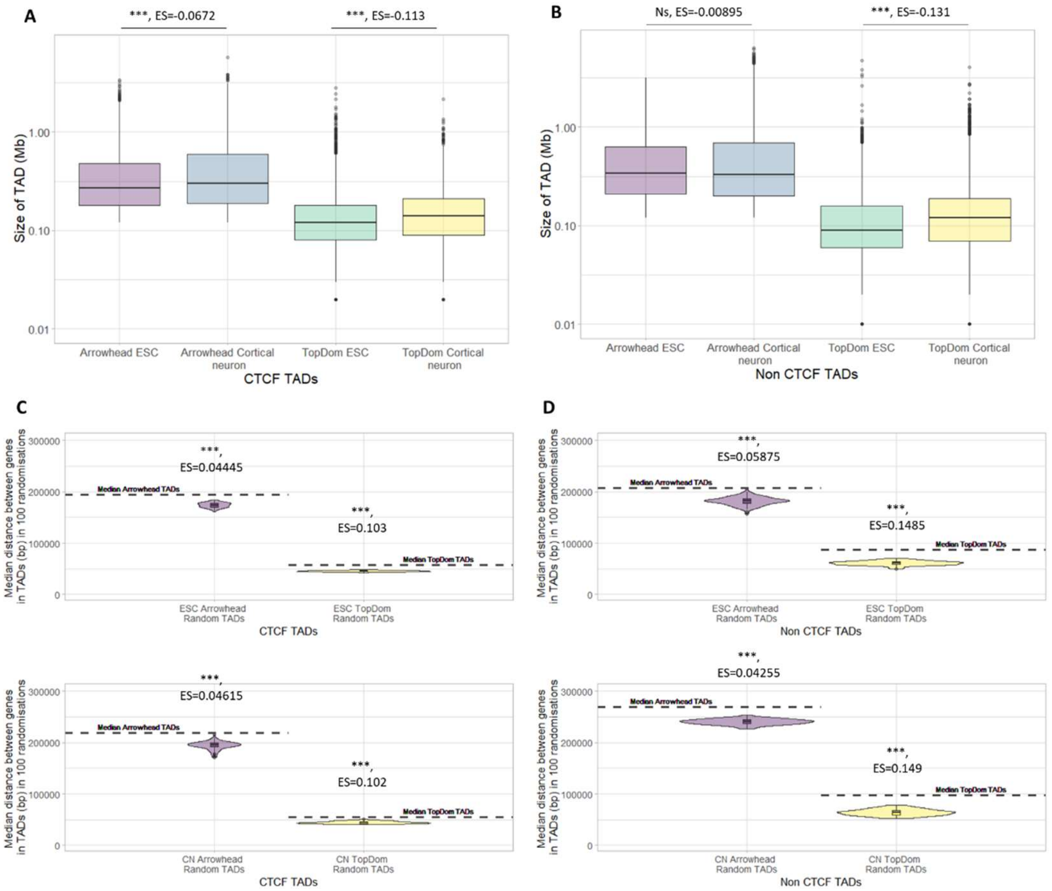


***Supplementary figure 3: Features of autosomal TADs split into CTCF TADs and non CTCF TADs.*** *A) Size of CTCF TADs called using Arrowhead and TopDom on ESC and cortical neuron Hi-C (plotted on a log10 scale) (Wilcoxon test, P-value: p<0.001 = ***, p<0.01 = **, p<0.05 = *, ES= Effect size calculated using r for Wilcoxon). B) Size of non CTCF TADs called using Arrowhead and TopDom on ESC and cortical neuron Hi-C (plotted on a log10 scale) (Wilcoxon test, P-value: p<0.001 = ***, p<0.01 = **, p<0.05 = *, NS= not significant, ES= Effect size calculated using r for Wilcoxon). C) Median distance between genes in CTCF TADs (dotted line) vs the median distance between genes in 100 sets of random TADs (plotted on a log10 scale). (Wilcoxon test, Median p-value: p<0.001 = ***, p<0.01 = **, p<0.05 = *, ES= Median effect size calculated using r for Wilcoxon). D) Median distance between genes in non CTCF TADs (dotted line) vs the median distance between genes in 100 sets of random TADs (plotted on a log10 scale). (Wilcoxon test, Median p-value: p<0.001 = ***, p<0.01 = **, p<0.05 = *, ES= Median effect size calculated using r for Wilcoxon).*

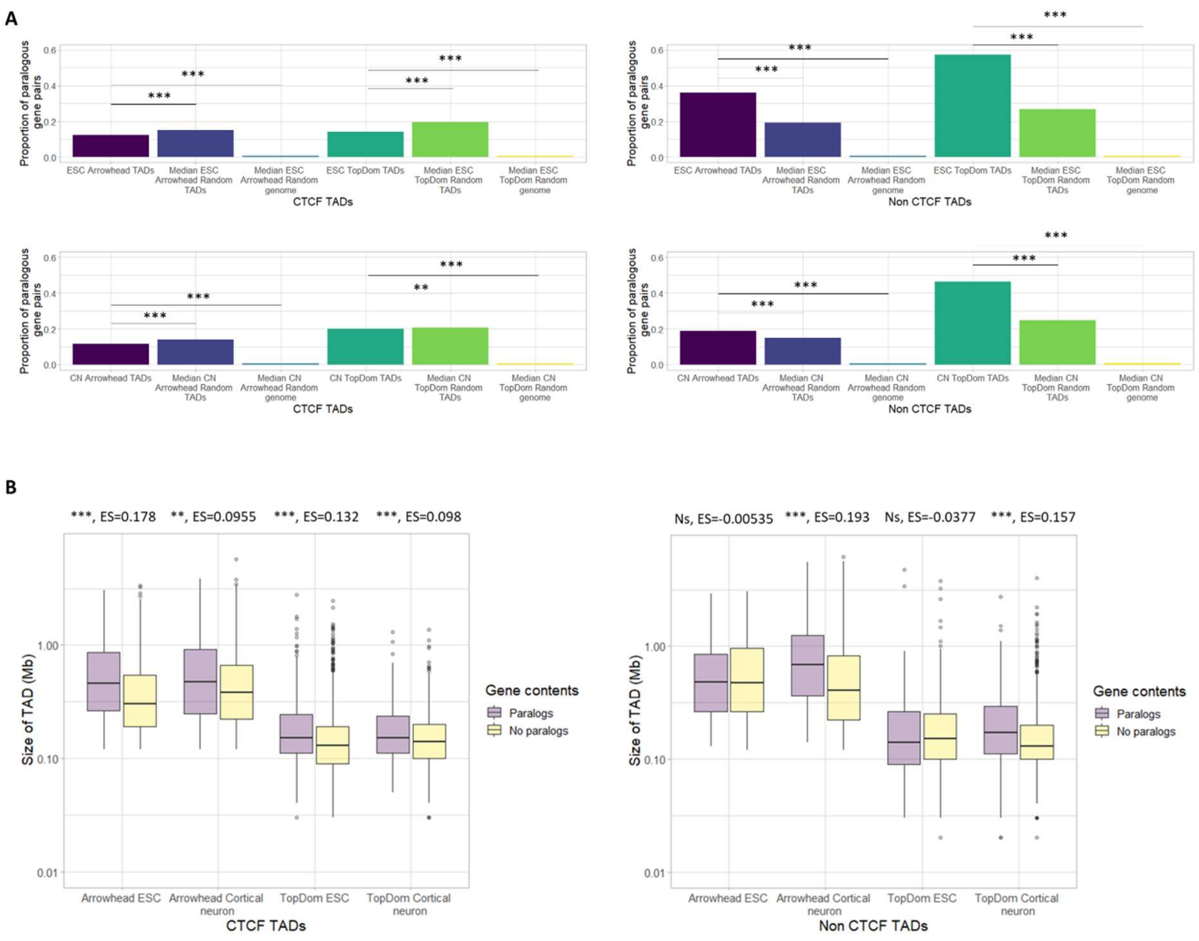


***Supplementary figure 4: Paralogous gene pairs in autosomal CTCF and non CTCF TADs.*** *A) Proportion of paralogous gene pairs in CTCF TADs (left) or non CTCF TADs (right) compared to the median proportion in 100 sets of random TADs, and the median proportion in 100 sets of random genome TADs in ESCs or cortical neurons (CN) (Fisher's exact test, Median p-value: p<0.001 = ***, p<0.01 = **, p<0.05 = *). B) Size of CTCF TADs (left) or non CTCF TADs (right) containing pairs of paralogs vs (with >1 gene) containing no pairs of paralogs (plotted on a log10 scale) (Wilcoxon test, p-value: p<0.001 = ***, p<0.01 = **, p<0.05 = *, NS= not significant, ES= Effect size calculated using r for Wilcoxon).*

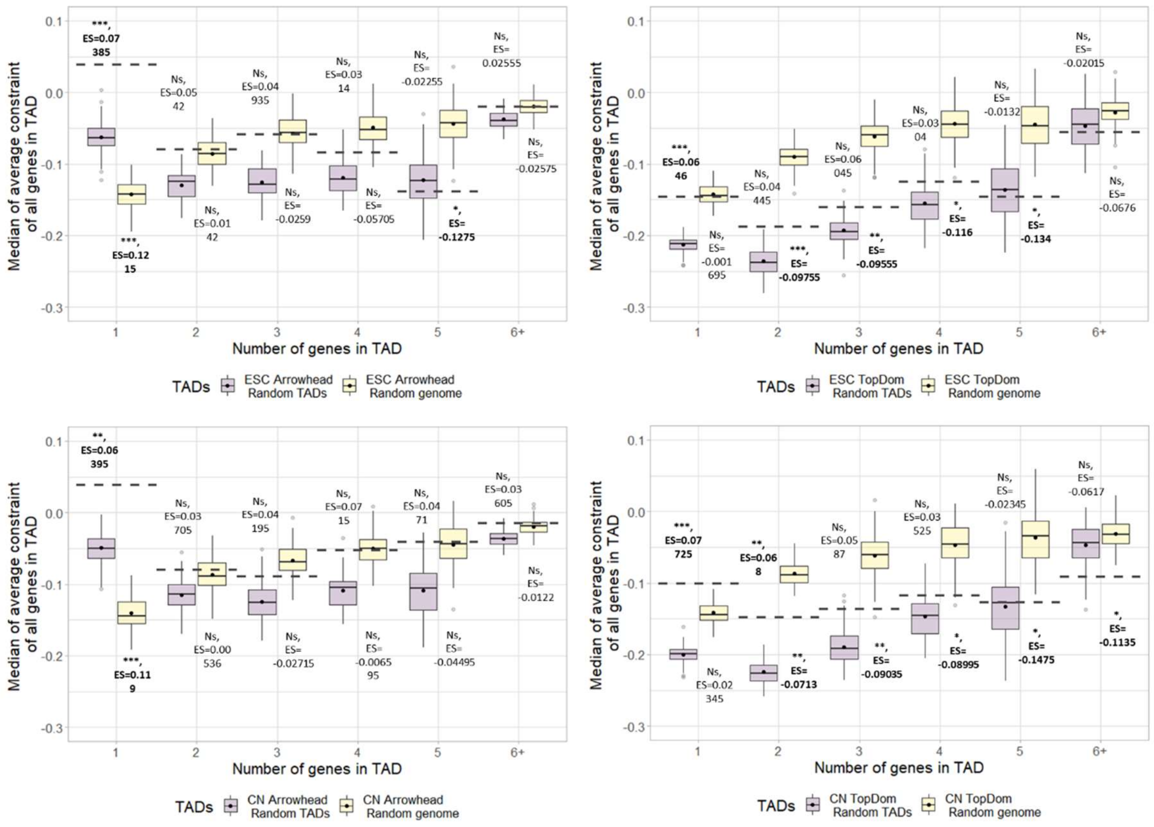


***Supplementary figure 5: Average constraint of genes in TADs vs random TADs/random genome autosomal TADs.*** *The average gene constraint score for all genes within a TAD/random TAD was calculated to create a score per TAD. TADs were binned based on the total number of genes they contained and the median score for each bin was plotted. The dotted line shows the median score in TADs and the distributions show the median scores in 100 sets each of: random TADs and random genome TADs. Dots indicate mean of the distribution (Wilcoxon test, FDR corrected median p-value (corrected for 6 tests, corresponding to the number of groups on the x axis): p<0.001 = ***, p<0.01 = **, p<0.05 = *, NS= not significant, Median ES= Effect size calculated using r). P-values shown above the bars for random TADs and below the bars for random genome TADs.*

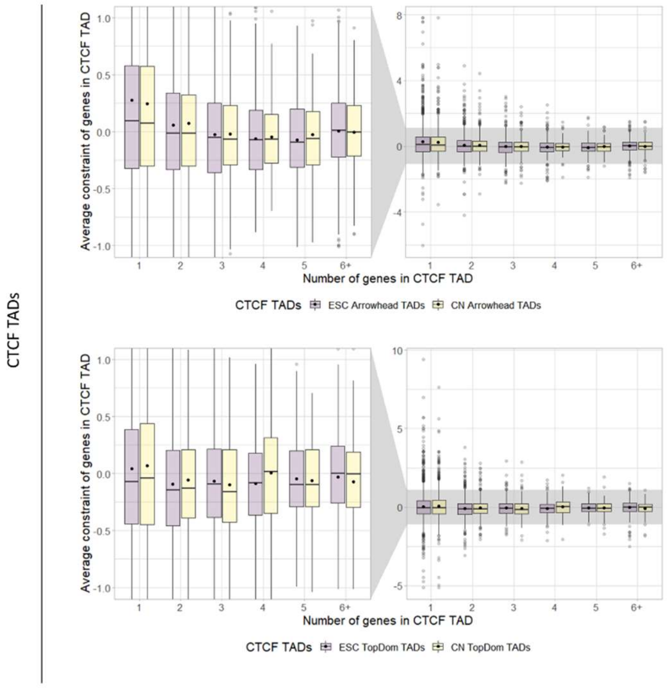


| Wilcoxon Rank Sum between average gene constraint in TADs containing 1, 2, 3, 4, 5, 6+ genes | | | | | |
|---|---|---|---|---|---|
| No. of genes in CTCF TAD | No. of genes in CTCF TAD | ESC Arrowhead CTCF TADs P value | CN Arrowhead CTCF TADs P value | ESC TopDom CTCF TADs P value | CN TopDom CTCF TADs P value |
| 1 | 2 | 0.00019066 | 0.0678011 | 0.01009089 | 0.2756724 |
| 1 | 3 | 2.3432E-06 | 0.0219328 | 0.37741675 | 0.2076122 |
| 1 | 4 | 1.1485E-05 | 0.0317589 | 0.43898017 | 0.812955 |
| 1 | 5 | 0.00019066 | 0.0678011 | 0.99871842 | 0.7150125 |
| 1 | 6+ | 0.00031607 | 0.0317589 | 0.37741675 | 0.7150125 |
| 2 | 3 | 0.17313556 | 0.4378897 | 0.37741675 | 0.4908926 |
| 2 | 4 | 0.14200985 | 0.5017333 | 0.43898017 | 0.3033182 |
| 2 | 5 | 0.17313556 | 0.5641502 | 0.28768686 | 0.6829552 |
| 2 | 6+ | 0.45950417 | 0.996844 | 0.00417874 | 0.4908926 |
| 3 | 4 | 0.78511764 | 0.996844 | 0.99871842 | 0.2274994 |
| 3 | 5 | 0.78511764 | 0.996844 | 0.43898017 | 0.3615419 |
| 3 | 6+ | 0.02401745 | 0.3360436 | 0.04427234 | 0.2768191 |
| 4 | 5 | 0.92836419 | 0.996844 | 0.47141251 | 0.6829552 |
| 4 | 6+ | 0.01155458 | 0.3360436 | 0.0850145 | 0.6829552 |
| 5 | 6+ | 0.02401745 | 0.4378897 | 0.37741675 | 0.7150125 |

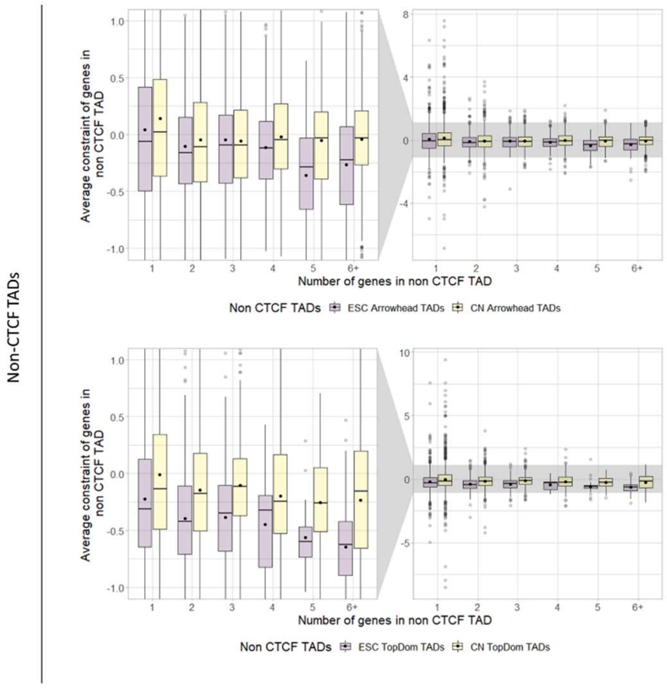


| Wilcoxon Rank Sum between average gene constraint in TADs containing 1, 2, 3, 4, 5, 6+ genes | | | | | |
|---|---|---|---|---|---|
| No. of genes in TAD | No. of genes in TAD | ESC Arrowhead TADs P value | CN Arrowhead TADs P value | ESC TopDom TADs P value | CN TopDom TADs P value |
| 1 | 2 | 0.072175909 | 0.001523437 | 0.035101111 | 0.06759245 |
| 1 | 3 | 0.576343748 | 0.003349508 | 0.153051217 | 0.70818333 |
| 1 | 4 | 0.339191053 | 0.142268378 | 0.186362822 | 0.11184923 |
| 1 | 5 | 0.000814325 | 0.042221313 | 0.008104076 | 0.11184923 |
| 1 | 6+ | 0.000814325 | 0.025211305 | 0.000218201 | 0.06759245 |
| 2 | 3 | 0.303837067 | 0.978673499 | 0.806809987 | 0.29776283 |
| 2 | 4 | 0.576343748 | 0.250614629 | 0.806809987 | 0.62184382 |
| 2 | 5 | 0.015851624 | 0.961334577 | 0.073118216 | 0.3881173 |
| 2 | 6+ | 0.072175909 | 0.086646626 | 0.013745163 | 0.46996673 |
| 3 | 4 | 0.576343748 | 0.255463445 | 0.806809987 | 0.18279625 |
| 3 | 5 | 0.002697125 | 0.961334577 | 0.063520948 | 0.11714742 |
| 3 | 6+ | 0.011455364 | 0.130417997 | 0.013745163 | 0.18279625 |
| 4 | 5 | 0.015003606 | 0.499635951 | 0.22296551 | 0.70818333 |
| 4 | 6+ | 0.072175909 | 0.978673499 | 0.140870203 | 0.79920265 |
| 5 | 6+ | 0.302168978 | 0.374334873 | 0.670402143 | 0.77792573 |

***Supplementary figure 6: Average constraint in CTCF autosomal TADs or non CTCF autosomal TADs binned by number of genes****. Distribution of mean constraint scores of all genes in CTCF TADs (top) or non CTCF TADs (bottom). Dots indicate mean of distribution. Tables showing FDR corrected p-values of differences between the groups in the graphs calculated with the Wilcoxon test. Significant p-values are highlighted red.*

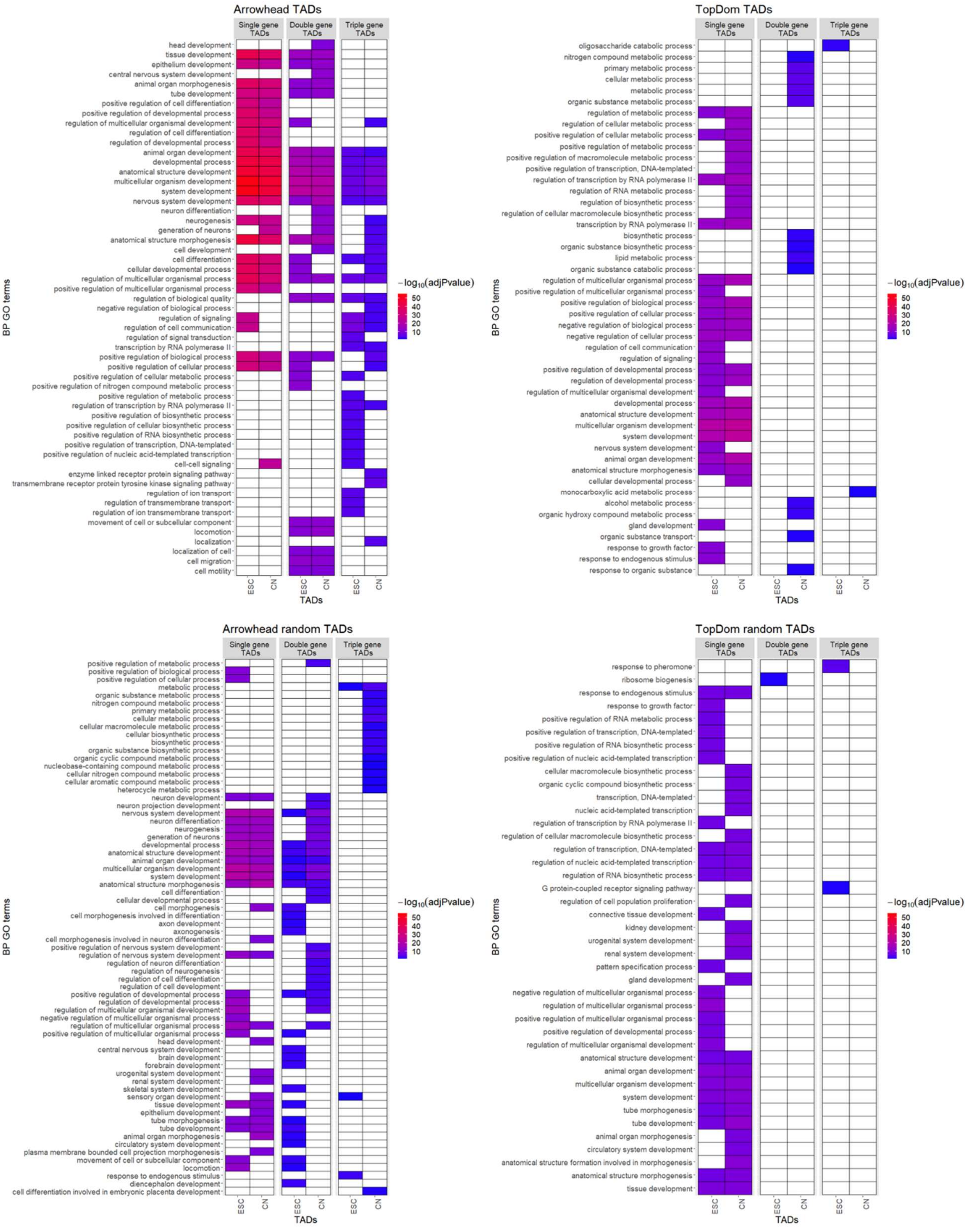


***Supplementary figure 7: Biological processes GO term functional enrichment of genes in TopDom TADs and Random TADs.*** *GO term enrichments calculated for genes within Arrowhead TADs,*

*TopDom TADs, one example of random Arrowhead TADs and one example of random TopDom TADs containing a single gene, two genes, or three genes. Only the top 25 most significant ($p<0.05$ multiple testing corrected using the “gSCS” option) GO terms are shown.*

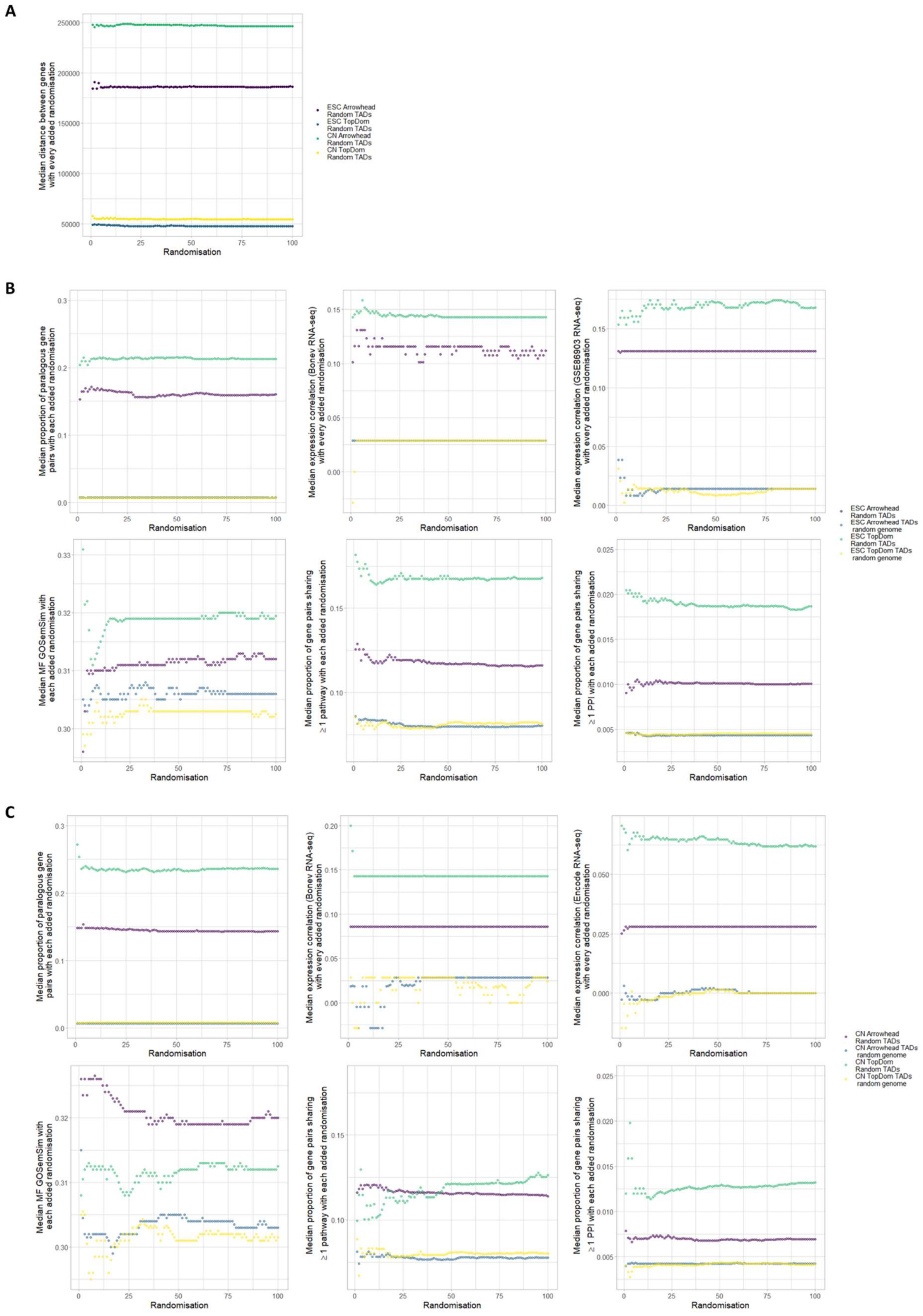
A
B
C
Median distance between genes with every added randomisation
Randomisation
ESC Arrowhead Random TADs
ESC TopDom Random TADs
CN Arrowhead Random TADs
CN TopDom Random TADs
Median proportion of paralogous gene pairs with each added randomisation
Median expression correlation (Bonev RNA-seq) with every added randomisation
Median expression correlation (GSE86903 RNA-seq) with every added randomisation
Median MF GOSemSim with each added randomisation
Median proportion of gene pairs sharing ≥ 1 pathway with each added randomisation
Median proportion of gene pairs sharing ≥ 1 PPI with each added randomisation
ESC Arrowhead Random TADs
ESC Arrowhead TADs random genome
ESC TopDom Random TADs
ESC TopDom TADs random genome
Median expression correlation (Encode RNA-seq) with every added randomisation
CN Arrowhead Random TADs
CN Arrowhead TADs random genome
CN TopDom Random TADs
CN TopDom TADs random genome

***Supplementary figure 8: Median value of measurers tested in random TADs with each added random TAD set:*** *For each measure the median begins to converge before 100 randomisations. A) Median distance between genes B) For ESC random TADs/random genome TADs: Median proportion of paralogous gene pairs, median expression correlation between gene pairs calculated using RNA-seq from Bonev et at. or GSE86903 et al., median MF GO semantic similarity between gene pairs, median proportion of pairs of genes sharing ≥1 pathway and median proportion of pairs of genes sharing ≥1 PPI. C) For cortical neuron random TADs/random genome TADs: Median proportion of paralogous gene pairs, median expression correlation between gene pairs calculated using RNA-seq from Bonev et at. or encode forebrain, median MF GO semantic similarity between gene pairs, median proportion of pairs of genes sharing ≥1 pathway and median proportion of pairs of genes sharing ≥1 PPI..*

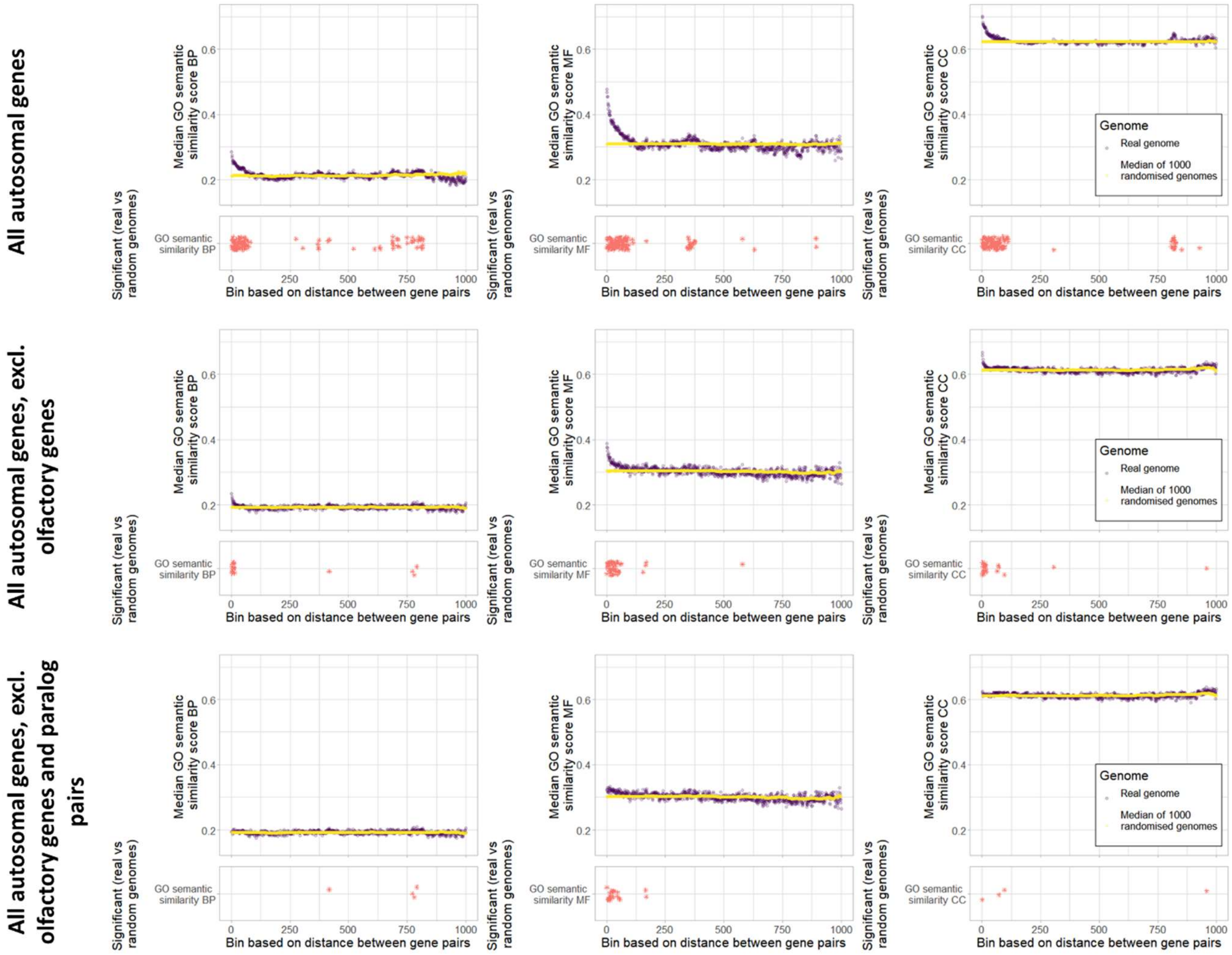


***Supplementary figure 9: Pairwise GO similar scores against genomic binned distance.*** *Distribution of GO semantic similarity for pairs of genes binned by distance in the real genome vs 1000 random genomes. In the top row of panels all autosomal genes are included, in the middle row olfactory genes are excluded and in the bottom row all olfactory genes are excluded along with any pairs of paralogs. Significance/bottoms panels: Stars indicate bins with a significantly higher GO semantic similarity in the real genome vs 1000 random genomes (FDR corrected p-value<0.05).*
